# Quantitative analysis of fibroblast migration reveals migratory states characterized by force generation, cell shape and motion

**DOI:** 10.64898/2026.05.06.723282

**Authors:** Elizabeth M. Davis, Max A. Hockenberry, Mitchell T. Butler, Harrison H. Truscott, Noah J. Shaul, James E. Bear, Timothy C. Elston

## Abstract

Cell migration depends on coordinating cell shape changes with force generation, yet how these processes are integrated remains unclear. Here, we combine live-cell imaging with traction force microscopy and computational analysis to quantify cell morphology, motility and force generation in migrating fibroblasts. We find that traction force magnitudes display a multimodal distribution, suggesting discrete migratory regimes. Using a Hidden Markov Model, we identify distinct force states that exhibit differences in shape and motion metrics, and show that individual cells transition between force states over time. To test the role of cytoskeletal organization in establishing the identified states, we analyzed cells lacking *Arpc*2, which disrupts branched actin assembly. Despite reduced forces and altered morphology, these cells also exhibit three migratory states. State transitions occur more frequently in cells lacking *Arpc2* and unlike normal cells their protrusion geometry is force dependent. Together, our findings show that cell migration is organized into discrete mechanical states that couple morphology, motility and force generation.

**SUMMARY STATEMENT:** Fibroblast motility involves distinct migratory states. These states exist independent of branched actin. However, state transition frequencies, traction force magnitudes and protrusion geometry are branched actin dependent.

## INTRODUCTION

Cell migration plays a fundamental role in many physiological processes, including development, morphogenesis, and immune function (Delgado and Lennon-Duménil, 2022; Friedl and Gilmour, 2009; SenGupta et al., 2021; Shellard and Mayor, 2021) and is often dysregulated in diseases such as cancer metastasis and autoimmune disorders. Cell migration emerges from coordinated cycles of membrane protrusion, adhesion formation and actomyosin contractility (Petrie and Yamada, 2015). In mesenchymal cells such as fibroblasts, these processes are prominently organized within the lamellipodium, a wide, sheet-like protrusion at the leading edge composed of a dense network of branched and linear actin filaments (SenGupta et al., 2021; Wu et al., 2012). Within this region, integrin-based adhesions form, mature and disassemble, linking the cytoskeleton to the extracellular matrix and enabling cells to sense and respond to mechanical cues (Alonso-Matilla et al., 2025). Through actomyosin contraction, cells transmit forces to their environment to drive forward movement (Weißenbruch and Mayor, 2024). Spatiotemporal coordination of protrusion, adhesion and retraction is required for effective cell motility. One challenge for elucidating the mechanisms that drive cell motility is that these processes occur over varying timescales: intracellular dynamics such as actin remodeling and adhesion turnover occur over seconds to minutes, whereas whole cell behaviors, such as migration, traction force generation, and shape change, emerge over minutes to hours (Brückner and Broedersz, 2024). Another complicating factor is that cell populations exhibit substantial heterogeneity, and ensemble-averaged measurements can obscure meaningful structure in the data (Altschuler and Wu, 2010; Huang, 2009). As a result, understanding cell migration requires approaches that capture changes in shape, motion, and force generation across both space and time.

Previous studies have examined pairwise relationships between motility, traction forces, and cell geometry (Lemmon and Romer, 2010; Messi et al., 2020; Rape et al., 2011; Tanimoto and Sano, 2014), but often lack temporal resolution or long-duration observations. A systematic approach that integrates all three features at the single-cell level over physiologically relevant timescales remains lacking. Therefore, we developed an integrated experimental and computational framework to study how cell shape, motion and traction forces measured at the scale of minutes are integrated over the scale of hours. A preliminary analysis based on quantitative metrics for cell motility suggested that migrating fibroblasts transition between distinct migratory states. To test this hypothesis, we applied stochastic modeling using hidden Markov models (HMMs), which have been applied previously in the prediction of cell morphological and motility states (Degerman et al., 2009; Gordonov et al., 2016; Held et al., 2010; Memmos et al., 2025; Mohammadi et al., 2022). Next, we took advantage of conditional knockout of the *Arpc2* gene to generate cells with defined changes in cell shape and motion (Methods). *Arpc2* null cells lack the ability to form branched actin and therefore cannot form lamellipodia (Rotty et al., 2015). These cells migrate slower than control cells, fail to migrate up gradients of extracellular matrix (haptotaxis), but are capable of chemotaxis and durotaxis (Hakeem et al., 2023; King et al., 2016; Wu et al., 2012). Thus, comparing wild-type and *Arpc*2 null cells allowed us to probe the relationships between cell shape, migration and traction force. Our results demonstrate fibroblast migration is organized into discrete, mechanically defined states that exist independent of branched actin networks. They also provide important insights into how cell shape and force generation are coupled to produce directed motion, laying the foundation for further investigations into cell migration.

## RESULTS

### Capturing the coordinated dynamics of cell shape, motility and traction forces across minutes to hours

To understand the relationship between cell shape, cell motion and traction forces, we imaged a GFP-expressing clonal mouse dermal fibroblast cell line (JR20) derived from an *Arpc*2 conditional knockout mouse (Methods). These cells were allowed to migrate on compliant PDMS substrates (∼20 kPa) functionalized with fluorescently labeled beads to allow for traction measurements. Using cytoplasmic GFP signal, we developed a computational platform that enabled automated cell outline segmentation and tracking of individual cells (Fig. 1A) as well as the calculation of metrics for traction force (Fig. 1B), motility (Fig. 1C) and shape (Fig. 1D). Cell motion was characterized using speed (total path length/time) and migration persistence (net displacement/total path length; D/T) (Fig 1C). To quantify cell shape, we computed a set of morphological metrics that have been successfully applied in many different contexts (Che et al., 2022; Chen et al., 2016; D’Orazio et al., 2020; Kenry, 2024; Vasilevich et al., 2020) ranging from predicting tumor growth and metastatic potential (Conner et al., 2024) to serving as a biological fingerprint for chondrocyte phenotype (Selig et al., 2023). Specifically, we used area as a measure of cell size, eccentricity to quantify deviations from circularity (where 0 indicates a circle and values closer to 1 indicate more oblong shapes), and solidity for boundary irregularity (where values closer to zero correspond to irregular morphologies) (Fig. 1D). Traction forces were computed from bead displacements (Fig. 1A) using the u-interforce Matlab software (Han et al., 2015). To characterize force generation, we computed the average traction force magnitude for each time frame (Fig. 1B). This extensive dataset allowed us to investigate how cell geometry, motility and traction forces are related over time scales relevant for cell physiological events such as migration. While previous work has explored some of these parameters in combination, evaluation of all of them simultaneously over many hours is necessary to dissect the contribution of each component and how they interrelate.

**Figure 1.**
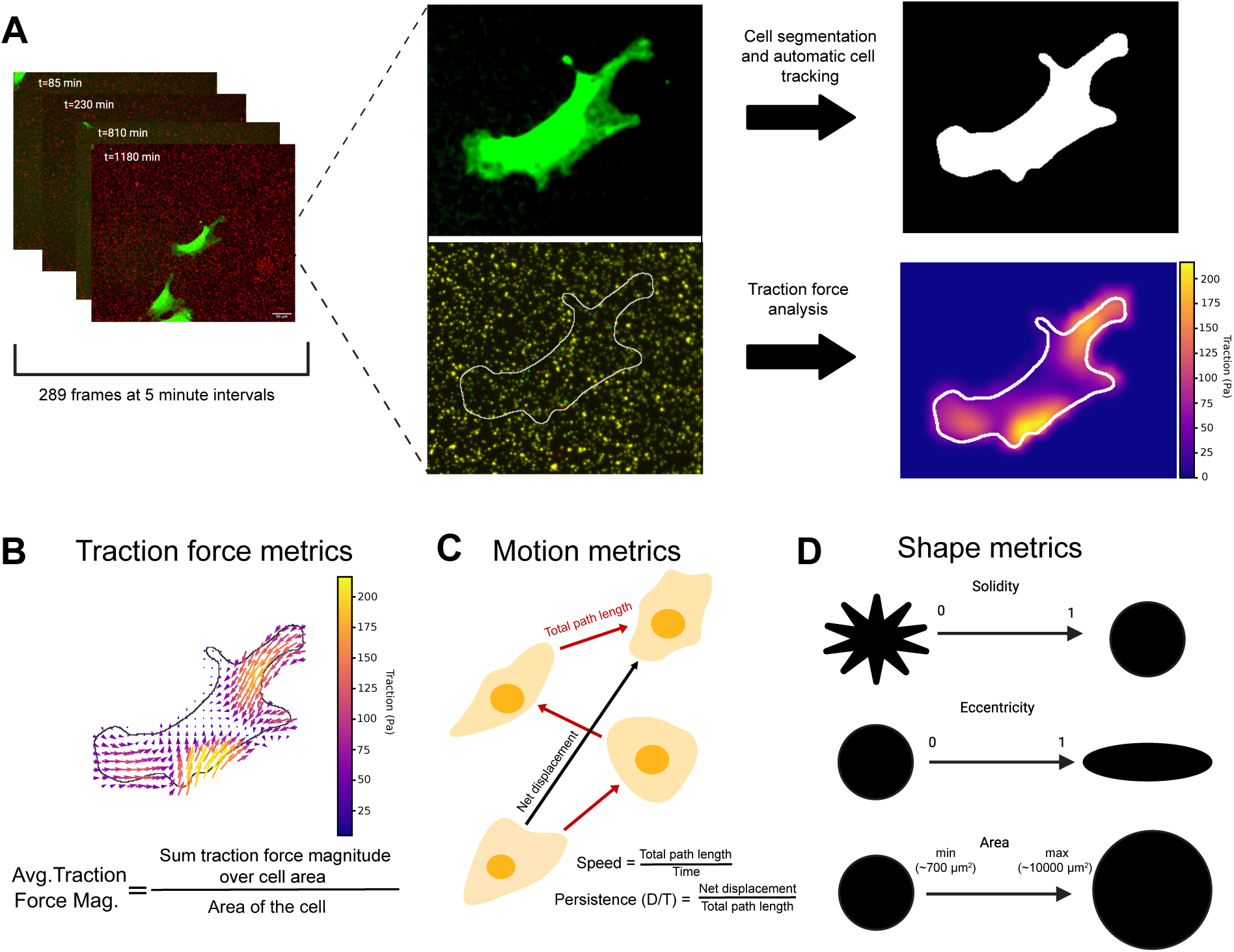
Development of a cell analysis pipeline to quantify traction force, motion and shape over time. (A) MEFs expressing cytoplasmic GFP were imaged at 5 minute intervals over the span of 24 hours on a compliant PDMS substrate coated with fluorescent polystyrene beads. From the cytoplasmic GFP, a binary mask of the entire cell was generated and the cell’s position was tracked over time using an automated pipeline. Cell induced displacements of the fluorescent beads were used to calculate traction forces generated by the cell. (B) The traction force vectors over the area of a cell were used to compute the average traction magnitude for each time point. (C) From the cell positions, motility metrics such as speed and persistence were calculated. (D) Using the cell masks, shape metrics such as solidity, eccentricity, and area were calculated.

We compared cell shape, motion, and force measurements between cells and summarized each metric by calculating its median value over the duration of the track. We then examined how these cell-level metrics covary by comparing pairwise combinations (Fig. 2A-H). Comparing shape and motility, we found that cell speed and area are negatively correlated, with smaller cells moving faster (Fig. 2A) and more persistently (Fig. 2B) than larger cells. However, there was not a statistically significant correlation when comparing solidity to speed and persistence (Fig. 2C, D). A comparison of force with motion metrics revealed that force per area inversely varies with speed and persistence (Fig. 2E, F), while a similar comparison of force and shape metrics demonstrated that large, smooth cells tend to generate more force per area than smaller, irregular shaped cells (Fig. 2G, H). Overall, our observations agree with previous work (Leal-Egaña et al., 2017). Interestingly, while solidity does not appear to be correlated with motility, it is correlated with average force generation, similar to cell area and motility-related metrics. This suggests a complex interplay between shape, motion, and traction force exists for migrating fibroblasts.

**Figure 2.**
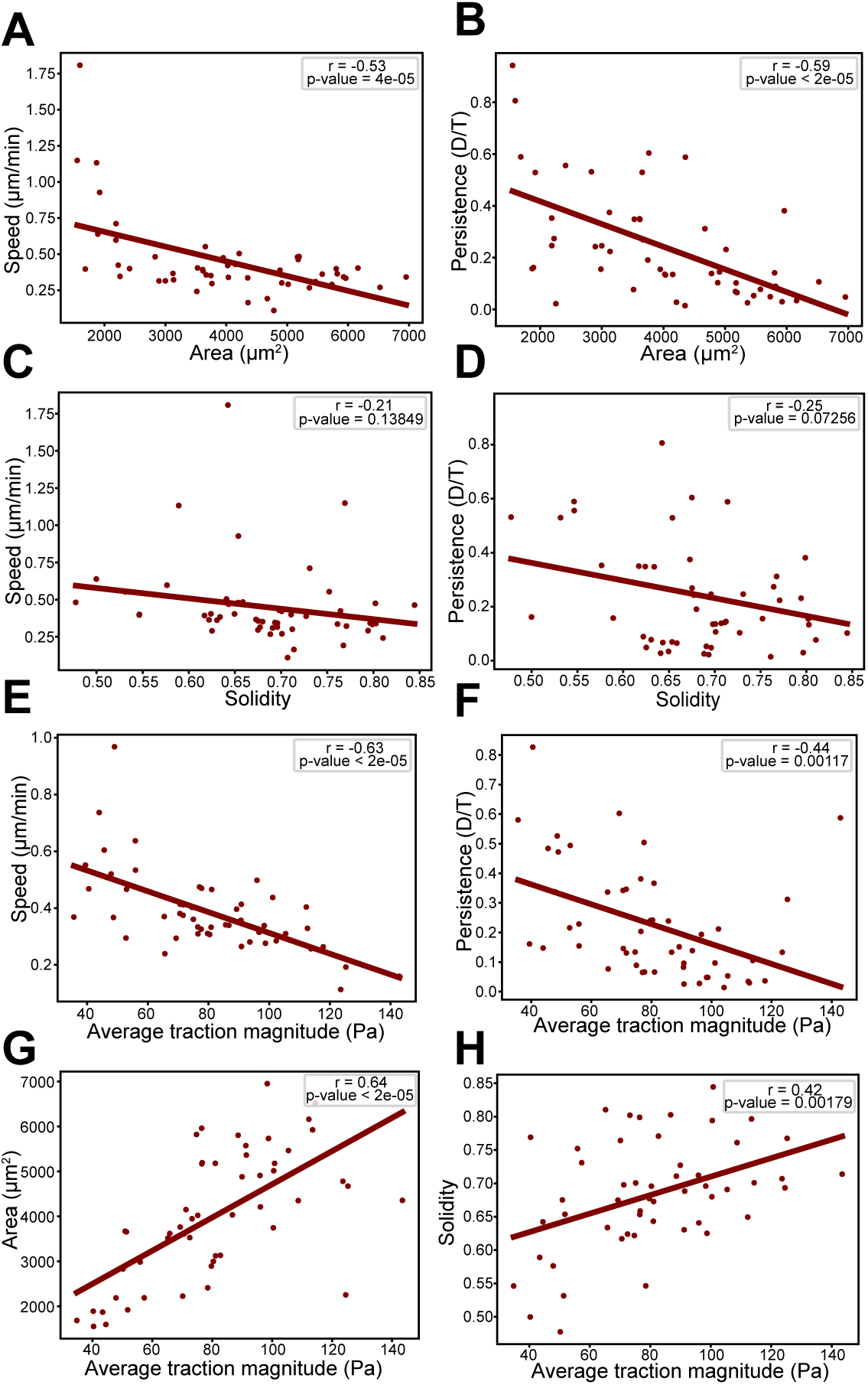
Correlation analysis of shape, motility, and force metrics. Scatter plots comparing metrics for shape and motility: (A) area and speed, (B) area and persistence, (C) solidity and speed, (D) solidity and persistence. Comparing metrics for force and motility: (E) average traction magnitude and speed, (F) average traction magnitude and persistence. Comparing metrics for force and shape: (G) average traction magnitude and area and (H) average traction magnitude and solidity. Each point on the plots represents the median value for one cell over the course of its track (N=52 cells). Linear regression was performed for each plot and the resulting line is plotted. The Pearson correlation coefficient and associated p-value are displayed on each plot. Statistical significance of the Pearson correlation coefficient is determined with a two-tailed t-test based on the t-distribution with n-2 degrees of freedom. A significance level of 0.05 was used.

### Traction force magnitude links cell shape and motility

One clue to understanding the observed relationships between force, shape and motion metrics is the observation that the distribution of traction force magnitudes exhibits a multimodal distribution (Fig. 3A). This suggests that subpopulations of cells representing distinct migratory states might be responsible for the observed behavior. To test this possibility and determine if individual cells transition between different migratory states, we turned to stochastic modeling. We employed a Hidden Markov Model (HMM) to test for distinct migratory states and gain insight into how traction forces are related to cell shape and motility. HMMs are designed to predict “hidden” states of a system from noisy time series data of experimentally measurable quantities that are functionally related to the hidden states. HMMs have been used in a wide variety of applications, including the prediction of cell morphological and motility states (Degerman et al., 2009; Gordonov et al., 2016; Held et al., 2010; Memmos et al., 2025; Mohammadi et al., 2022). We sought to determine if applying a HMM model to the traction force data would support the existence of subpopulations, and if so, do relationships between shape and motility exist within these subpopulations that are masked when the full data set is considered. To conduct our analysis, we used a three-state Gaussian HMM (Rabiner, 1989). Our rationale for selecting a 3-state model was based on a comparison of models with varying number of states ranging from 1-10 (Methods). The 3-state model accurately captures the traction force distribution (Fig. 3A) while not overfitting the data (Fig. S3). The three states correspond to a low force state centered around 50 *Pa*, a medium force state centered around 80 *Pa* and a high force state centered around 119 *Pa*. The HMM allows us to predict the state of the system at each time point. A representative example of a cell undergoing three transitions during imaging is shown in Fig. 3B with the time series for the average traction force magnitude and associated probability of being in each force state over time shown in Fig. 3C. Of the 52 cells analyzed, around 75% of the cells underwent a state transition over the duration of its trajectory with an average transition rate of 0.077 ± 0.011 transitions per hour (Fig. S6). This allowed us to compute transition rates between the states. We found similar average transition rates to and from the low and medium force states and from the medium force state to high force state (Fig. 3D). Transitions from the high force state to the medium force state were highest and no transitions from the high to low force were observed. The average time spent in the low force state was around 5.6 hours, time spent in the medium force state was around 9.6 hours, and the high force state was around 5.9 hours (Fig. S6).

**Figure 3.**
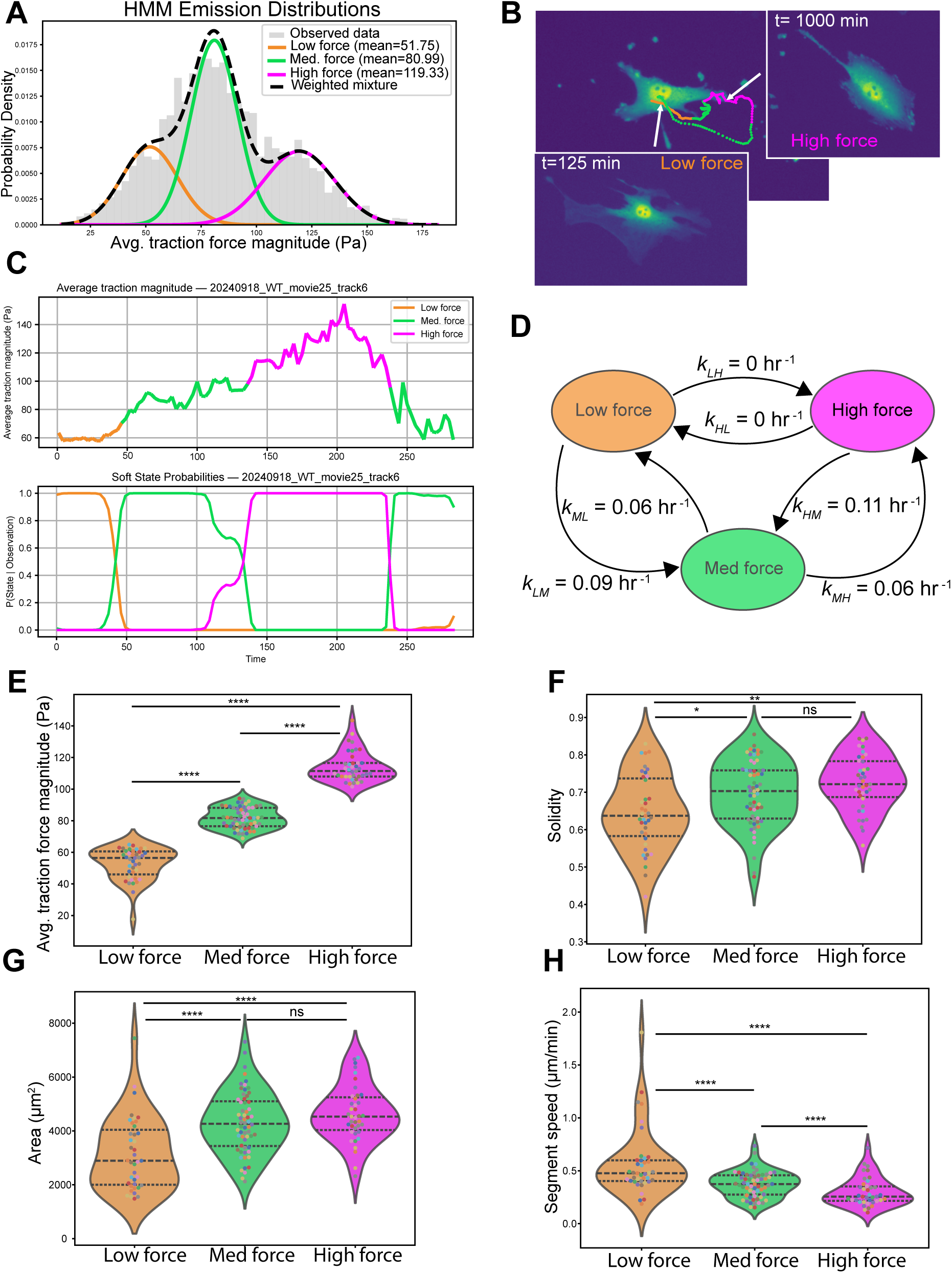
Hidden Markov model predicts migratory states with distinct motility and shape features. (A) Predicted traction force states from the hidden Markov model. (B) Example of a cell that undergoes three transitions between states. The cell begins in the low force state, transitions to the medium force state, transitions to the high force state, and then returns of the medium force state. (C) Time series for the average traction force magnitude showing the predicted states (top panel) and probability of being in each state (bottom panel) for the cell in (B). (D) Transition rates between the three states obtained from the model. (E-H) Comparison between states of average traction magnitude, solidity, area, and speed. Each dot represents a continuous track segment defined by the migratory state and dot colors correspond to individual cells (N=52 cells with an average of 2.65 segments per cell track). Black dotted lines display the quartiles. Statistical significance was determined using a permutation test (see Methods). P-values ≤ 0.05, ≤ 0.01, ≤ 0.001, and ≤ 0.0001 are represented as *, **, *** and ****, respectively.

We next compared how force, motion and shape metrics differed between the predicted migratory states. To do this we computed the median values of the metrics over track segments in which cells remained in the same state. First, we compared the average traction force magnitude for the predicted states (Fig. 3E). As expected, the states showed significant differences in the median values of the average traction forces. Interestingly, the migratory states also correlated with observed differences in shape and motion parameters. Cells in the low force state have lower solidity (more irregular shape) than those in the high force state (Fig. 3F) and are smaller (Fig. 3G). Cells in the low force state tend to move faster than those in the high force state (Fig. 3H). To establish the statistical significance of the observed differences between the 3 states, we performed permutation tests to compute p-values for the state-dependent differences (Methods). We found significant differences between the low and high force states for all measured parameters. These observed differences in shape and motility between the predicted states suggest that cellular geometry influences cells’ ability to generate forces and translocate. To further probe this relationship, we genetically perturbed the actin cytoskeleton, hypothesizing that causing changes in actin network geometry would affect shape, motility and force generation.

### *Arpc2* KO results in decreased solidity, slower speed and increased persistence

A key cellular component that underlies motility and cell morphology is the actin cytoskeleton. The Arp2/3 complex is responsible for nucleating branched actin which structures lamellopodia (Fig. 4A) (Goley and Welch, 2006; Rotty et al., 2013), sheet-like protrusions that form at the leading edge of migrating cells and are sites of nascent adhesion formation and force generation (Choi et al., 2008). Therefore, to probe how force, shape and motility are affected by actin network geometry, we abrogated the Arp2/3 complex function by the genetic deletion of the *Arpc*2 gene, encoding a critical subunit in the Arp2/3 complex, which results in cells losing branched actin and thus lamellopodia (Fig. 4B) (Rotty et al., 2015). The *Arpc2* KO cells are visually distinct from WT cells, with KO cells having a spikier appearance (Fig. S6). This change in appearance is quantified by solidity, with the *Arpc2* KO cells having lower values of solidity than WT cells (Fig. 4C). *Arpc2* KO cells are also more elongated (Fig. 4D) and generally smaller (Fig. 4E) than WT cells. Even in the absence of branched actin, *Arpc2* KO cells are still able to migrate. However, when compared to WT cells, *Arpc2* KO cells move slower (Fig. 4F) and more persistently (Fig. 4G), consistent with previous findings (Hakeem et al., 2023). Despite moving more slowly than WT cells, *Arpc2* KO cells generate lower average traction magnitudes than WT cells (Fig. 4H). This observation is surprising because WT cells exhibited a negative correlation between force and speed, so one might expect *Arpc2* KO cells to move faster than WT cells. This discrepancy suggests that a different migratory strategy may be employed by *Arpc2* KO cells.

**Figure 4.**
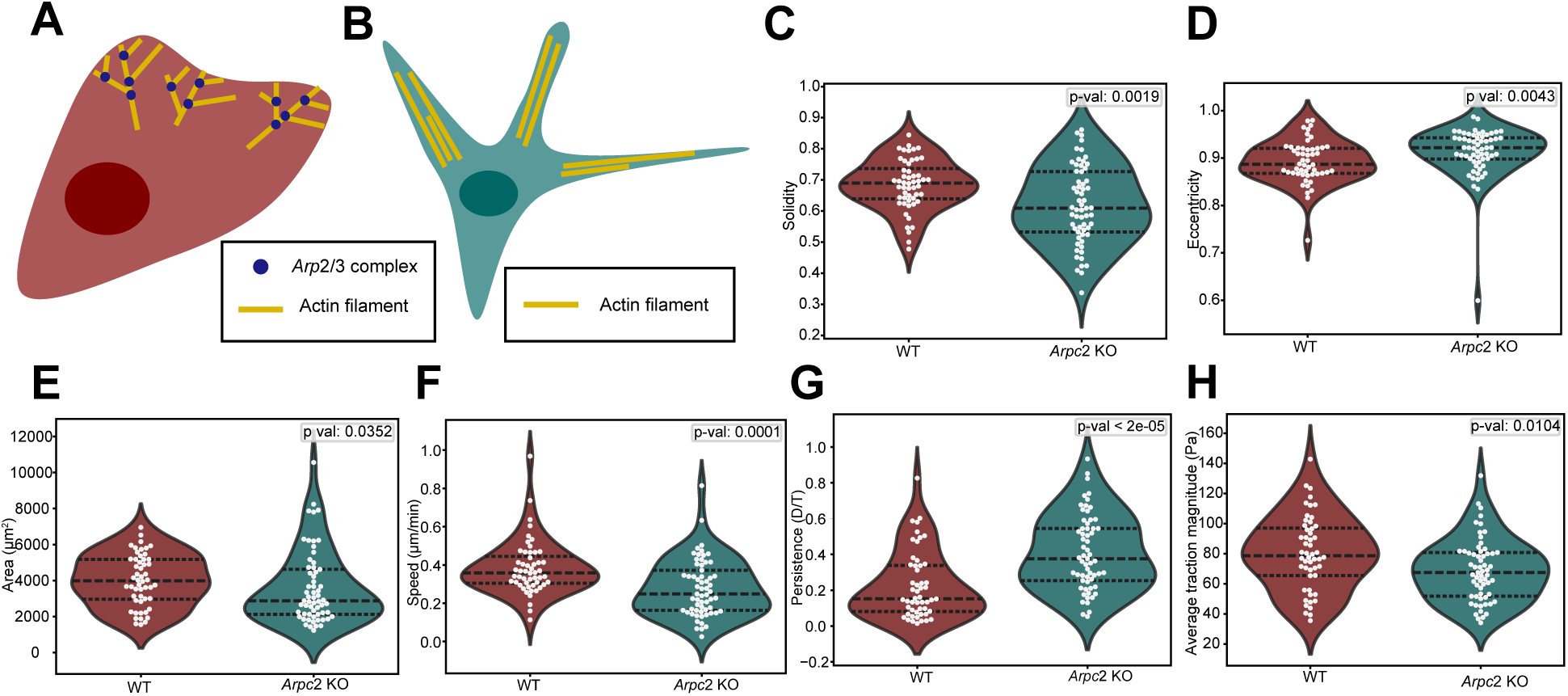
Comparison of *Arpc*2 KO cells and WT cells. (A) A WT cell can form lamellipodia due to the *Arp*2/3 complex, which nucleates branched actin. (B) An *Arpc*2 KO cell lacks the *Arp*2/3 complex so branched actin cannot form, resulting in a bundled actin morphology with long, thin protrusions. (C-H) Comparison of median track values for solidity, eccentricity, area, speed, persistence and average traction magnitude between WT and *Arpc*2 KO cells. Each dot represents the median metric value of one cell over the entire course of its track (WT: N=52 cells, *Arpc*2 KO: N=61 cells). Black dotted lines display the quartiles. Statistical significance was determined using the Mann-Whitney U test.

### *Arpc2* KO cells show distinct shape, motion, and traction force relationships

For *Arpc2* KO cells, area and speed are negatively correlated (Fig. S4), similar to the relationship seen in WT cells but stronger. However, unlike WT cells, which did not display any correlations between motion and solidity, *Arpc2* KO cells with higher solidity tend to move slower and more persistently than more irregularly shaped cells (Fig. 5A, B). We did not see a correlation between persistence and average traction magnitude (Fig. 5D) in contrast to WT cells. However, cells with a lower average traction magnitude tend to move faster, similar to WT cells (Fig. 5C). This similarity is surprising since *Arpc2* KO cells generate lower traction forces yet move slower than WT cells (Fig. 4F, H). This suggests that in *Arpc2* KO cells traction forces play a different role in generating cell motility. Finally, similar to WT cells, both area (Fig. 5E) and solidity (Fig. 5F) are positively correlated with average traction magnitude, but with slightly weaker correlation than WT cells. While WT and *Arpc2* KO cells present differences in shape, motion, and traction force between the two cell types (Fig. 4C-H), *Arpc2* KO cells still display a trimodal distribution of average traction force magnitude, similar to WT cells, once again suggesting the presence of subpopulations (Fig. 6A). Therefore, we fit a HMM to these data to test for three states and probe for relationships between shape and motion metrics within the states.

**Figure 5.**
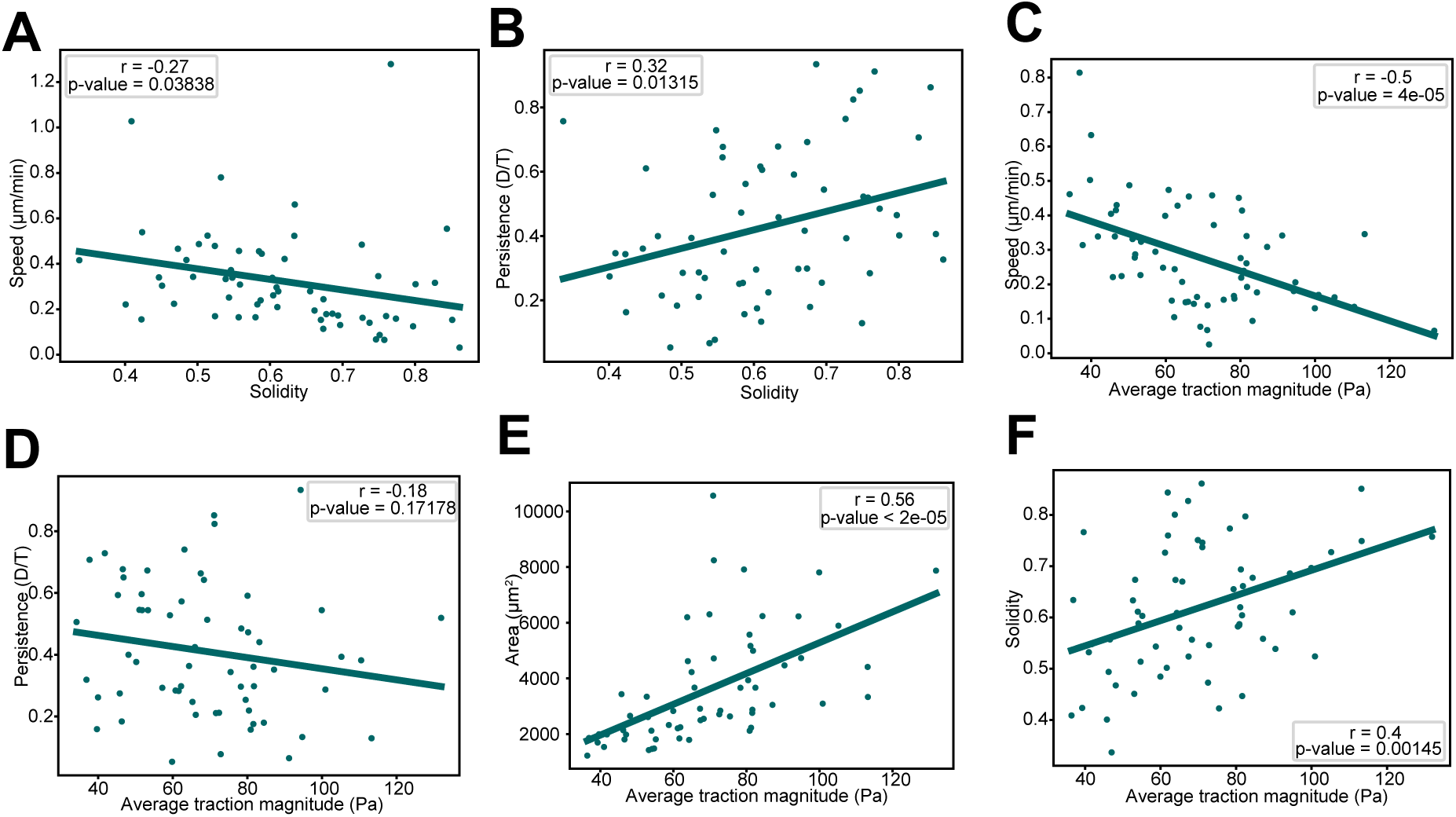
Correlation analysis of *Arpc*2 KO cells. (A-F) Scatter plots comparing shape (area and solidity), motion (speed and persistence) and average traction magnitude for *Arpc*2 KO cells. All plots except (D) show statistically significant correlations. Each point on the plot represents the median value for one cell over the course of its track (N=61 cells). Statistical significance of the Pearson correlation coefficient was determined with a two-tailed t-test based on the t-distribution with n-2 degrees of freedom. A significance level of 0.05 was used.

**Figure 6.**
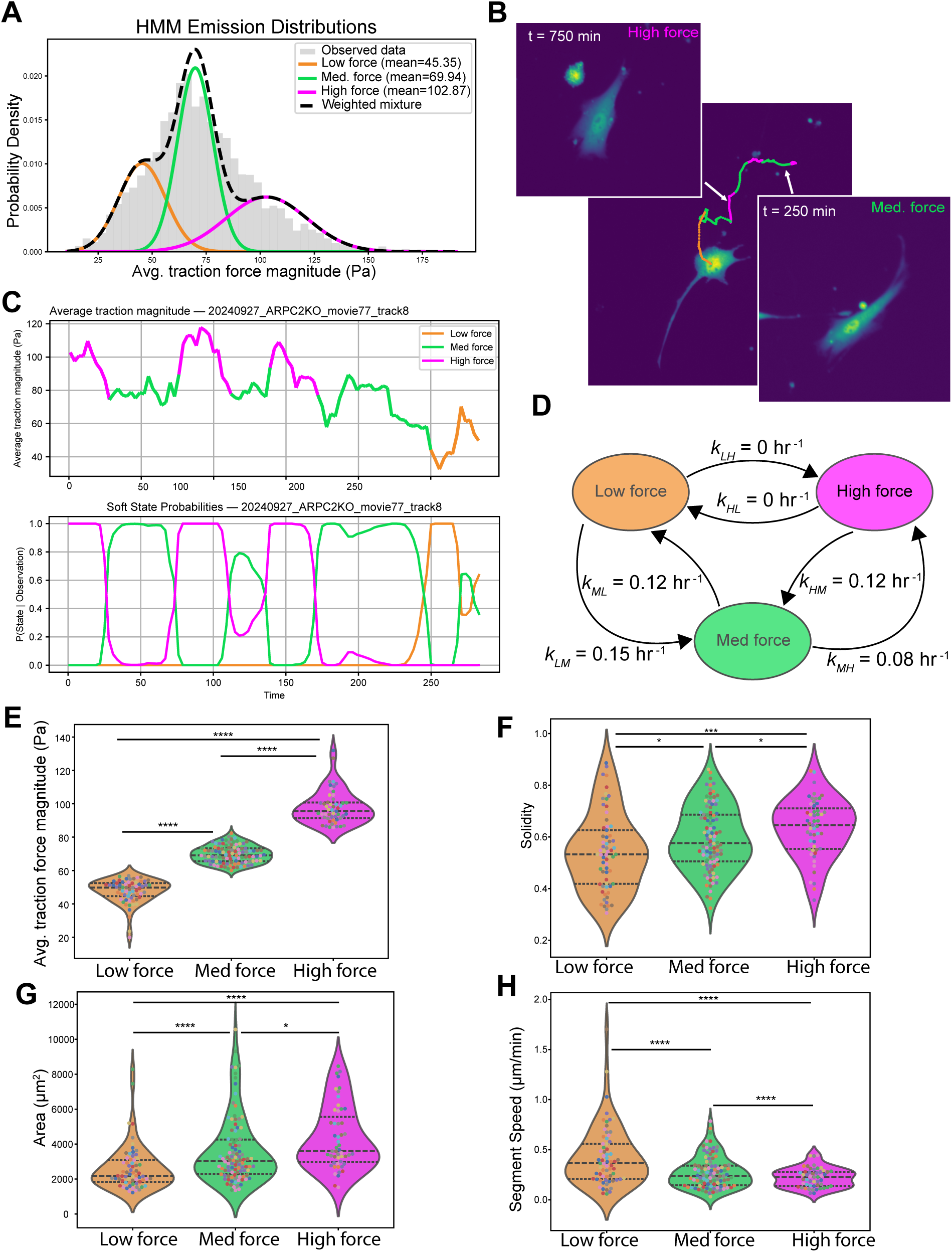
Hidden Markov model predicts migratory states with distinct motility and shape features for *Arpc*2 KO cells. (A) Predicted traction force states from the HMM for *Arpc*2 KO. (B) Example of a cell that undergoes 6 transitions between states. (C) Time series for the average traction force magnitude showing the predicted states (top panel) and probability of being in each state (bottom panel) for the cell in (B). (D) Transition rates between the three states obtained from the HMM. (E-H) Comparison between states of average traction magnitude, solidity, area, and speed. Each dot represents a continuous track segment defined by the migratory state and dot colors correspond to individual cells (N=61 cells with an average of 3.49 segments per cell). Black dotted lines display the quartiles. Statistical significance was determined using a permutation test (see Methods). P-values ≤ 0.05, ≤ 0.01, ≤ 0.001, and ≤ 0.0001 are represented as *, **, *** and ****, respectively.

### Migratory states persist in *Arpc2* KO cells with altered transition dynamics

Similar to the WT data, the traction force distribution for *Arpc2* KO cells was well captured by a three-state Gaussian HMM (Fig. 6A) (Methods). For these cells, the low force population was centered at 45 *Pa*, the medium force population was centered at 70 *Pa* and a high force population was centered at 103 *Pa* (Fig. 6A). Overall, the states of the *Arpc2* KO are shifted to lower forces and closer together than the WT results. A representative example of an *Arpc2* KO cell that undergoes six transitions during imaging is shown in Fig. 6B with the time series for the average traction force magnitude and associated probability of being in each force state over time shown in Fig. 6C. *Arpc2* KO cells have faster transition rates overall, with the largest increases in the transitions between the low and medium force states (Fig. 6D). Once again, no transitions occur between the low and high force states. Of the 61 *Arpc2* KO cells analyzed, 79% of the cells underwent transitions over their trajectory with an average transition rate of 0.12 ± 0.012 transitions per hour (Fig. S6) which is higher than we observed with WT cells. Despite undergoing more transitions, *Arpc2* KO cells spent a similar portion of time in each state compared to WT cells (Fig. S6). This suggests that cells lacking Arp2/3 have more dynamic traction forces.

As expected, the average traction magnitude calculated from cell track segments predicted to be in the low, medium and high force states are well separated (Fig. 6E). Similar to WT cells, there are differences in shape and motion between the three states. *Arpc2* KO cells have highest solidity in the high force state indicating a more convex shape (Fig. 6F). Additionally, cells are on average largest in the high force state (Fig. 6G). Like WT cells, *Arpc2* KO cells move slowest in the high force state (Fig. 6H), and there is a difference in persistence between the three states (Fig. S7) with cells in the high force state moving the least persistently. Once again, we performed permutation tests to establish the statistical significance of these results (Methods). The states predicted from the HMM for *Arpc2* KO cells had similar differences in shape and motion between the states as the HMM for WT cells despite their different protrusion structures. This leads us to investigate how protrusions vary between these two cell types and force states.

### Cells lacking branched actin display distinct protrusion characteristics and less dynamic cell boundaries

Morphological features that differ between WT and *Arpc2* KO is the appearance of their protrusions (*Arpc2* KO cells appear to have longer and thinner protrusions) and the dynamics of the cell boundaries (WT cells appear to change shape more frequently). To quantify differences in protrusion geometry, we identified protrusions as regions of the membrane that extend beyond the cell body and measured their width and length (Fig. 7A) (Methods). Comparing protrusion geometry between WT and *Arpc2* KO cells, we found a difference in protrusion number, width and length with *Arpc2* KO cells producing fewer, narrower and longer protrusions (Fig. 7B-D). Next, we measured protrusive and retractive activity along the boundary of the cell membrane using the ProActive module from the computational platform CellGeo (Tsygankov et al., 2014). To compute protrusive and retractive activity, we compared the area between the cell boundary in successive frames (Fig. 7E). First we summed the areas for protrusions and retractions and normalized this number by the total cell area to compute total boundary activity as a percent of total cell area. WT cells displayed more total boundary activity compared to *Arpc2* KO cells (Fig. 7F) and this was also true when protrusion and retraction activity were considered separately (Fig. S9).

**Figure 7.**
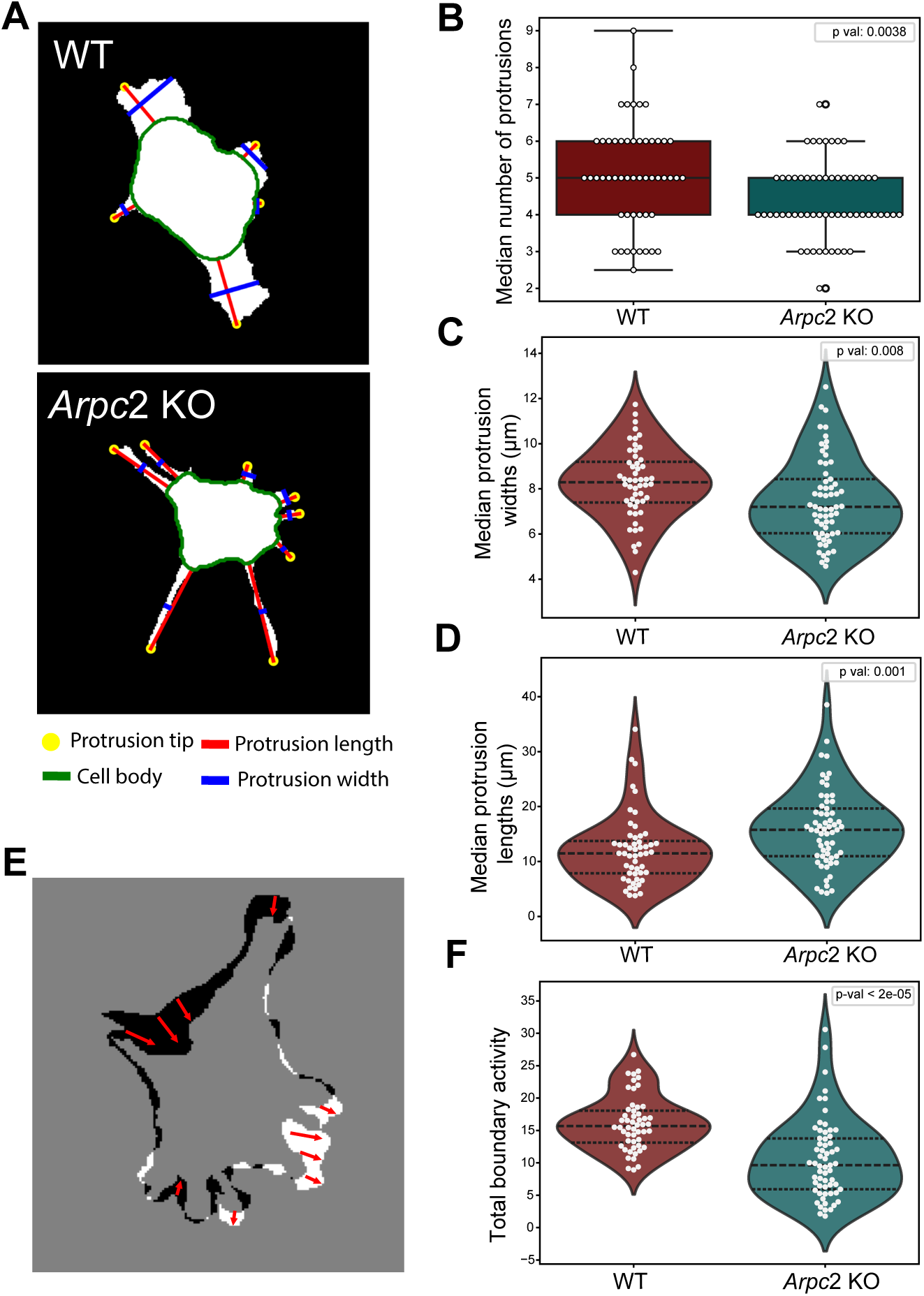
Comparison of protrusions between WT *Arpc*2 KO cells. (A) Protrusions were defined as regions outside the cell body (green line). The length (red line) and width at the midpoint of the protrusion (blue line) were calculated. (B) Median number of protrusions over a cell’s track for WT and *Arpc*2 KO cells. (C) Median per cell protrusion widths for WT and *Arpc*2 KO cells. (D) Median per cell protrusion lengths for WT and *Arpc*2 KO cells. (E) Schematic illustrating boundary activity. The cell boundaries for consecutive time points (t_i_, t_i+1_) are compared. Regions where the boundary protrudes are indicated in white and regions where it retracts are indicated in black. (F) Median per cell boundary activity. Boundary activity was computed as the sum of protrusion and retraction area divided by the average cell area at t_i_ and t_i+1_. (WT: N=52 cells, *Arpc*2 KO: N=61 cells). Black dotted lines display the quartiles. Statistical significance is determined with the Mann-Whitney U test.

Because *Arpc2* KO cells have a lower average traction force magnitude than WT cells, we wondered if differences existed between protrusions in the predicted HMM force states. The number of protrusions was state dependent for WT cells with more protrusions generated in the high force state than the low force state (Fig. 8A, left panel). The same trend appeared to hold for *Arpc2* KO cells (Fig. 8A, right panel). However, this difference did not reach the set threshold for statistical significance. We found that protrusion width did not vary by state for WT cells, but *Arpc2* KO cells produced narrower protrusions in the low force state (Fig. 8B). There were no differences in peripheral protrusion length between force states (Fig. S10) for either WT or *Arpc2* KO cells. Finally, we examined boundary activity within the predicted force states from the HMM. We found that protrusive activity depended on force state, with lower force states displaying higher protrusive activity than higher force states for both WT and *Arpc2* KO cells (Fig. 8C). While there were also significant decreases in retractive activity with increasing force state for both cell types (Fig. S10), it was less evident than protrusive activity.

**Figure 8.**
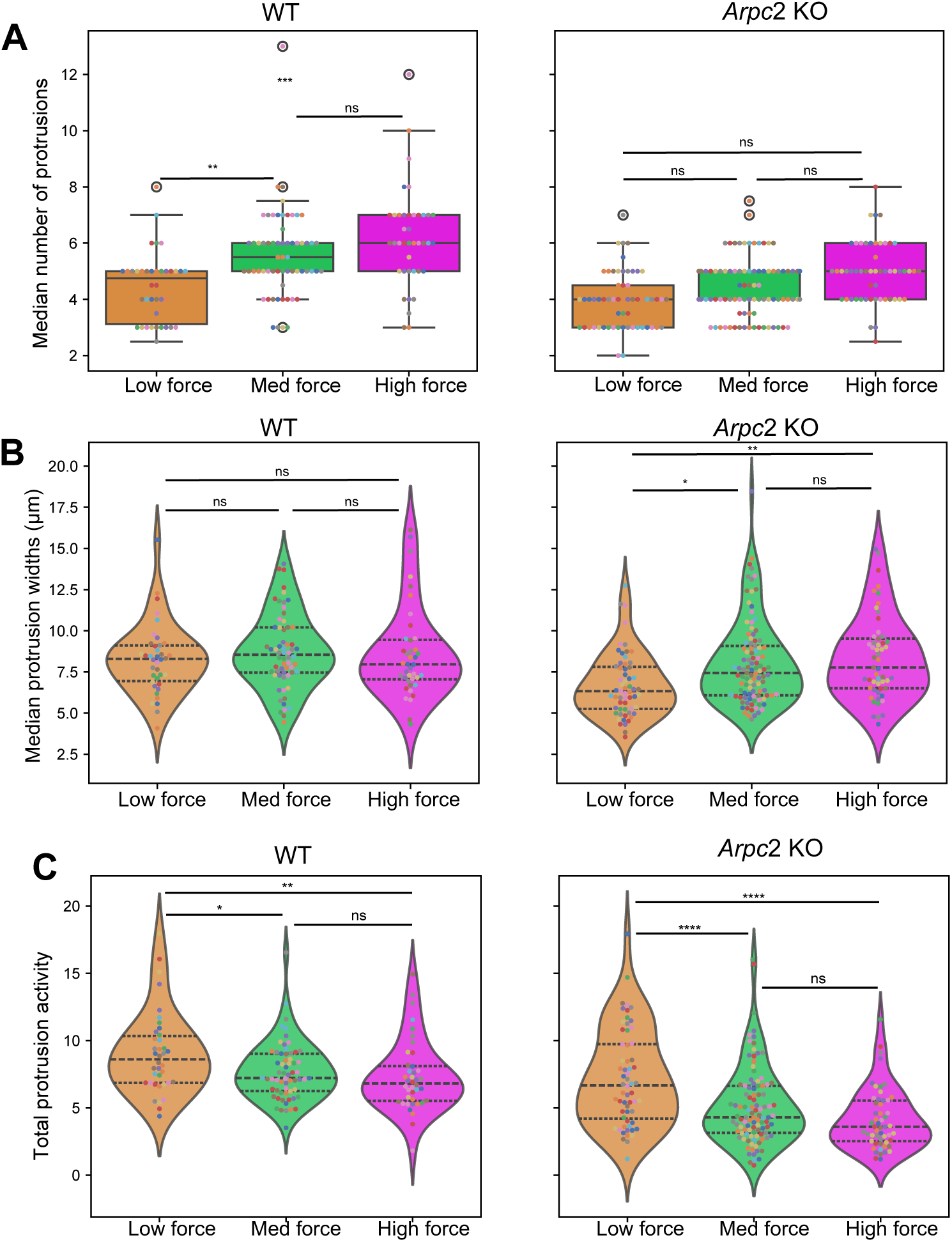
Comparison of protrusions between migratory states. (A) Median of the median number of protrusions per track segment for each migratory state for WT and *Arpc*2 KO cells. (B) Median protrusion widths per track segment for each migratory state for WT and *Arpc*2 KO cells. (C) Median boundary activity per track segment for each migratory state for WT and *Arpc*2 KO cells. (WT: N=52 cells, *Arpc*2 KO: N=61 cells). Black dotted lines display the quartiles. Statistical significance is determined with a permutation test using a significance level of 0.05 (Methods). P-values ≤ 0.05, ≤ 0.01, ≤ 0.001, and ≤ 0.0001 are represented as *, **, *** and ****, respectively.

## DISCUSSION

In this work, we demonstrated that random fibroblast migration occurs in three discrete traction force states. Using Hidden Markov Modeling applied to time series for traction force measurements, we inferred the state of each cell over time, and found that the migratory states exist on the time scale of hours and cells within each state displayed distinct shape and motility characteristics. *Arpc2* KO cells, which lack lamellapodia, display altered shape, motility, and force generation also exhibited three migratory states. Both cell types displayed increasing solidity and area and decreasing speed as force increased. However, *Arpc*2 KO cells displayed higher transition rates between the force states than WT cells, possibly reflecting the structural differences in the protrusions formed in each cell type (lamellipodia for WT; filopodia for *Arpc2* KO).

Analysis of protrusion geometry revealed that protrusion width correlates with traction force generation in *Arpc2* KO cells, suggesting that these protrusions are mechanically sensitive. This finding is consistent with previous work showing that *Arpc2* KO cells retain the ability to undergo durotaxis (Hakeem et al., 2023), suggesting that mechanically responsive peripheral protrusions may partially compensate for the loss of lamellipodia. We also found that *Arpc2* KO cells displayed lower levels of boundary activity compared to WT cells. However within the predicted HMM states both cell types displayed decreasing protrusive activity with increasing force state. This seems to suggest that while loss of the Arp2/3 complex reduces protrusive activity, it does not fundamentally affect protrusive mechanical coupling.

We found that cells in the low force state had a more irregular membrane boundary, were smaller and moved faster while cells in the high force state had a smoother membrane boundary, were larger and moved slower. Additionally, no transitions were observed between the low to high force states, thus requiring that cells visit the intermediate state when transitioning out of these states. The long dwell times observed within each state suggest that these transitions reflect stable migratory regimes rather than stochastic fluctuations. While our HMM captures more course-grained properties such as overall cell shape, it may be possible that there are finer-scale dynamics within each state. For instance, it has been shown that adhesion forces fluctuate over the scale of nanoseconds to mediate the sensing of extracellular matrix rigidity (Plotnikov et al., 2012). Previous work has shown that traction forces are spatially organized, increasing toward the cell periphery, where elevated stress triggers retraction, which occurs over the span of minutes (Messi et al., 2020). Although our state transitions occur on longer time scales than these edge dynamics, it is possible similar spatial patterns persist within each state, where each force state contains distinct sub-states that reflect cycles of protrusion and retraction. Investigating these spatial traction patterns within states could reveal whether each migratory regime is composed of unique internal dynamics or shared protrusion-retraction behaviors.

Our results suggest that changes between migratory states reflect underlying changes in the mechanical coupling between cell spreading, adhesion maturation and speed. As cells spread and adhesions mature, the overall traction force exerted increases. However, the strong adhesion to the extracellular matrix can also impede movement, thus highlighting a tradeoff between contractility and translocation. This tradeoff is captured by the motor-clutch model (Bangasser et al., 2013) which describes how myosin motors pulling on actin filaments transmit force to the substrate through stochastic clutch engagement of adhesions. Within this framework, our observed three state model could be interpreted in terms of different clutch engagement regimes. For instance, the low force state may correspond to a low clutch engagement regime characterized by low traction magnitude and irregular cell shape morphology, whereas the high force state may reflect a high clutch engagement regime with greater traction forces and larger and more evenly spread cells and slower migration. Interestingly, *Arpc*2 KO cells display higher transition rates (Fig. 6A) despite spending a similar amount of time in each state as WT cells (Fig. S6). This indicates that loss of the Arp2/3 complex does not alter the accessibility of migratory states, but instead affects the kinetics of transitions between them. Previous work has shown that focal adhesions in *Arpc*2 KO cells form at the base of the narrow protrusions and mature rapidly, in contrast to WT cells whose focal adhesions mature in the distal to proximal direction at a slower rate (Wu et al., 2012). Within the motor-clutch framework, this behavior could suggest more rapid cycles of clutch engagement and failure dynamics resulting from the altered actin architecture in the *Arpc*2 KO cells. Such changes may also introduce spatial differences in clutch binding and unbinding dynamics. These results suggest that branched actin architecture contributes to the temporal stability of traction force regimes during cell migration.

Broadly, our work shows that cell migration can be viewed in terms of transitions between discrete mechanical regimes that couple cell morphology and traction force generation. While previous studies have examined relationships between cell shape, motility and traction force over shorter time scales, our results have shown that these processes can give rise to stable phenotypic states over longer time scales. These state based behaviors provide a framework for understanding how feedback between cytoskeletal organization, adhesion dynamics and force transmission governs migratory behavior. Future work integrating traction force measurements with focal adhesion and actin imaging will be important in linking these state transitions to their underlying molecular mechanisms. Additionally, probing how extracellular stiffness modulates transitions between migratory states may provide insight into how cells tune their mechanical behavior in different environments, with potential implications for processes such as tumor metastasis, where dynamic switching between migratory modes is thought to play a key role.

## Acknowledgements

We thank Saygin Gulec for his helpful advice on cell segmentation. We also thank Samuel Ramirez for his guidance in the computational pipeline in the early stages of this project.

## MATERIALS AND METHODS

### Loxless GFP cell line generation

In order for GFP expression to remain intact following Cre recombination at the Arpc2 locus, the lentiviral pLL5.0-GFP plasmid LoxP sites were removed from by amplifying the DNA sequences between them using primer sets 5’-acctagtgagcATTAAGGGTTCCAAGCTTAAGCGGC-3’ with 5’-cttaaacatgtcactcgTAGGTCCCTCGACGAATTGCCT-3’ and 5’-ggaacccttaatgctcacTAGGTCCCTCGACCTGCTGGAA-3’ with 5’-ACCTAcgagtgACATGTTTAAGGGTTCCGGTTCCACT-3’ before joining the two resulting PCR products via Gibson Assembly. Lentiviral production was performed using standard procedures. In brief, HEK293T cells were transfected with 500 ng each of the “loxless” pLL5.0-GFP, pCMV-VSV-G, pRSV-Rev, and pMDL-g/p plasmids using Xtremegene HP transfection reagent (Roche). Lentivirus was harvested 72 hours later and used to transduce a clonal Arpc2 conditional knockout mouse dermal fibroblast line (JR20) isolated from a previously described bulk population (Rotty et al., 2017) for 72 hours while supplemented with 4 ug/ml Polybrene. GFP-positive JR20 cells with moderate expression levels were purified via FACS.

### Microscopy

To create a robust, long time scale dataset to investigate random motility behavior, cytoplasmic GFP labelled mouse embryonic fibroblasts (MEFs) plated on PDMS substrates with a Young’s modulus of 20 kPa coated with Thermo fluorescent carboxylated polystyrene beads were imaged at 5 minute intervals over the course of 24 hours. These experiments yielded 84 10x movies of WT cells and 70 movies of *Arpc2* KO cells with an average of around 20 cells per movie. However, after enforcing strict filtering criteria of cells that were isolated and motile upon visual inspection, we were left with 52 movies of individual WT cells spanning 3 independent experiments and 61 movies of individual *Arpc2* KO cells spanning 2 independent experiments with most tracks around 16-24 hours in length (Fig. S1).

### Image processing

Both the GFP and RFP channels of the images undergo drift correction. This is accomplished by comparing the bead positions of each frame to the last frame of each movie since the last position contains the bead positions not under the influence of the cell’s applied forces. The displacements needed to shift each image for maximal alignment are found using the chi2_shift() function from the image_registration library (image_registration developers, 2023). This function uses the discrete Fourier transform to calculate the cross correlation to compare peaks between frames to determine the shift. The GFP images were used to generate binary masks of the cell cytoplasm using a combination of Cellpose (Stringer et al., 2021), with the pretrained cyto2 model using a diameter of 100 pixels, and Fouier high pass filtering. Using the binary masks, the custom automated analysis pipeline written in Python removes cells that exceed or are less than a specified size. Additionally, cells were filtered to remove cells that leave the field of view.

### Tracking, shape, and motion metric calculation

After removing cells that did not meet size requirements, exited the field of view, or were identified as artifacts, the remaining cells were tracked across frames. Cell centers were determined using the approximate medioid method, which is computed by finding the point within the cell mask that minimizes the Euclidean distance to the median *x* and *y* coordinates of all pixels in the mask. Using these centers, we implemented a nearest neighbor tracking algorithm in which cells in consecutive frames were associated based on minimum Euclidean distance (Rosebrock, 2018). To reduce tracking errors, constraints such as maximum allowable displacement between frames and the maximum number of frames a cell could be absent before being assigned a new label were imposed. New labels were also assigned when cells newly appeared in the field of view.

For each resulting track, image data were cropped to a smaller field of view such that the entire track is contained within. This cropping was applied to GFP images, mask images, and bead images, reducing images from around 2000×2000 pixels to at most 400×400 pixels. This step reduced computational cost for downstream traction force calculations by excluding unnecessary background regions. Additionally, cell shape metrics such as area, convex area, eccentricity, orientation, perimeter, equivalent diameter, solidity, extent, major_axis length, and minor axis length were computed using the region_props() function from the skimage.measure library (Van Der Walt et al., 2014). Solidity was defined as the ratio of the cell area to its convex hull area, where values around zero denote a spikier, more irregular shape while a value of 1 is a perfectly solid, convex shape. Eccentricity was calculated by fitting an ellipse to the cell shape and is defined as the ratio of the focal distance to the major axis length, providing a measure of elongation. Area was computed as the number of pixels within the cell mask scaled by pixel area.

To quantify cell motion, centroid positions from each track were used to compute migration metrics. Due to the noise in centroid positions introduced fluctuations in cell shape, trajectories were smoothed using a non-overlapping moving average with a window size of three frames. Step sizes and turning angles as well as overall track metrics like speed and persistence were then calculated from the smoothed trajectories. Speed was defined as the total path length divided by elapsed time. Persistence was defined as the ratio of net displacement to total path length, where a value of 1 indicates perfectly directed motion and values approaching 0 indicate more diffusive movement.

### Traction force analysis

Traction forces were calculated from the movies of cells displacing the beads on the PDMS substrate using the u-interforce Matlab software (Han et al., 2015). Within the software, we used particle image velocimetry (PIV) to calculate the bead displacements with a template size of 21 pixels. To calculate the traction forces, we selected the Fourier transform traction cytometry (FTTC) method with a regularization parameter of 0.00001(King et al., 2022). The resulting tractions were saved as csv files to analyze them in our shape, motion, and traction force analysis code written in Python. Within our Python analysis script, we calculated traction force metrics such as average traction magnitude and dipole ratio. Average traction magnitude is the sum of the magnitude of the traction forces over the area of the cell divided by the area of the cell. The dipole ratio is obtained by finding the eigenvalues of the force dipole matrix. The ratio is therefore the normalized minor eigenvalue over the normalized major eigenvalue.

### Gaussian hidden markov model

Hidden markov models (HMMs) are used to infer states based on a set of measurable parameters. We used average traction magnitude as our measurable parameter for the HMM in order to predict migratory states. We implemented the hidden markov model using the hmmlearn Python package (hmmlearn developers, 2010). We use a HMM with Gaussian emissions since our distribution of average traction magnitude seemed to suggest a mixed Gaussian model. To initialize the means and covariances for the model, we begin by fitting a Gaussian mixture model with the selected number of hidden states in our HMM using the curve_fit() function from scipy.optimize (Virtanen et al., 2020). We trained for 100 iterations using the default convergence threshold of 0.01. We calculated the Akaike information criterion (AIC), Bayesian information criterion (BIC), and log likelihood (LL) to help inform us what the appropriate number of hidden states we should utilize. Following the elbow method we determined 3 states would be the appropriate number to use (Fig. S3A, Fig. S7A).

### Protrusion measurement

We defined the cell body by applying a morphological opening operation (erosion followed by dilation) to binary cell masks using skimage.morphology.opening() (Soille, 2004), an approach commonly used for removing protrusive structures (Barry et al., 2015; Hansel et al., 2019). To set the spatial scale of this operation, we used a disk-shaped structuring element with a radius proportional to cell size. Specifically, for each cell we computed the equivalent radius of a circle with the same area, 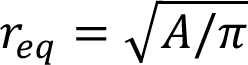, and defined the structuring element radius as a fraction of 𝑟_eq_. It was found that a ratio of 0.5 of the equivalent radius empirically produced the most accurate separation of cell body and peripheral protrusions. However, in some cases (particularly for elongated thin cell shapes), this radius resulted in no detectable cell body. To address this, we implemented an iterative method in which the structuring element radius was progressively reduced until a cell body was identified or a minimum radius threshold was reached. Additionally, for elongated cells, the opening operation occasionally produced two disconnected regions at opposite ends of the cell. To counteract this, we computed the convex hull of the resulting regions to obtain a single contiguous representation of the cell body. Peripheral protrusions were defined as the regions remaining after subtracting the cell body from the original mask, with only regions exceeding a minimum area threshold retained. For each peripheral protrusion, length was defined as the linear distance between the farthest point on the peripheral protrusion to the closest point on the cell body. Width was measured at the midpoint of this length by constructing a line perpendicular to the length axis and projecting nearby pixels onto this line. Examples of peripheral protrusions identified with this method are shown in Fig. S8.

### Statistical analysis

Statistical analyses for the Pearson correlation coefficient and the Mann-Whitney U test were calculated using the stats package from Scipy. To compare differences between the states predicted from the Hidden Markov Model, we performed a permutation test. This involved shuffling the data for the parameter being tested while maintaining the predicted HMM state labels to simulate the null hypothesis of there being no meaningful difference between the states. This process is repeated with 10,000 permutations of the shuffled data and for each permutation a test statistic is calculated. We chose to use the difference in median between groups as our test statistic. Once this process is completed, we have a null sampling distribution of the test statistic. We then compare the test statistic of the original, unshuffled data with the null sampling distribution and determine the proportion of permutations that have test statistics that are equal to or less than or greater than the observed value to give us our p-value. We represent p-values ≤ 0.05, ≤ 0.01, ≤ 0.001, and ≤ 0.0001 as *, **, *** and ****, respectively.

---

**Supplemental Figure 1.**
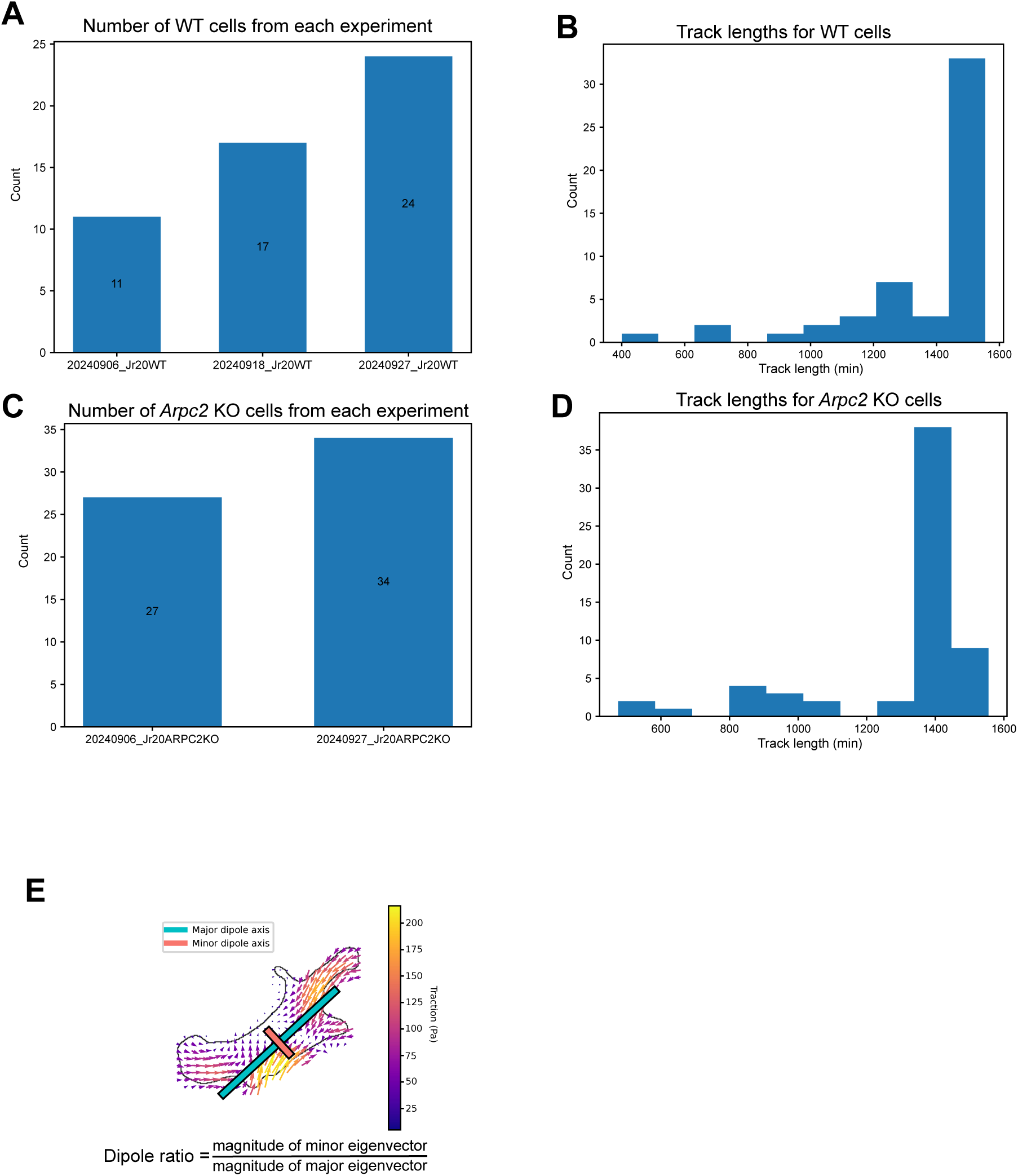
(**A**) Number of WT cells from each independent experiment. (**B**) Histogram showing the track lengths for all the WT cells. (**C**) Number of *Arpc*2 KO cells from each independent experiment. (**D**) Histogram showing the track lengths for all the *Arpc*2 KO cells. (**E**) Diagram displaying the force dipole on an example cell.

**Supplemental Figure 2.**
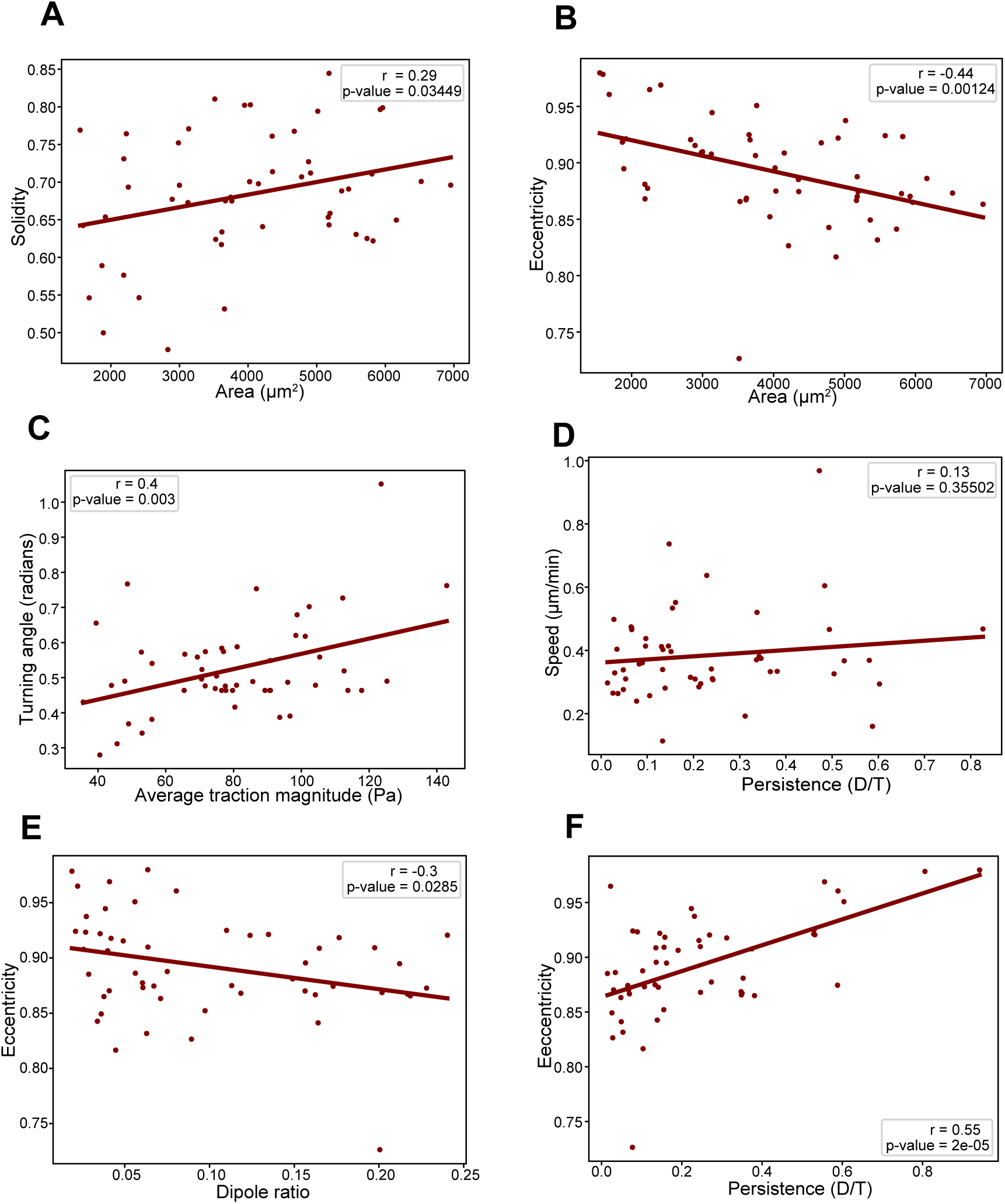
(**A-F**) Scatter plots comparing shape, motility and traction force metrics for WT cells. Each point on the plot represents the median value for one cell over the course of its track (N=52 cells). Linear regression was performed for each plot and the resulting line of best fit is plotted. The Pearson correlation coefficient and associated p-value is displayed on each plot. Statistical significance of the Pearson correlation coefficient is determined with a two-tailed t-test based on the t-distribution with n-2 degrees of freedom. A significance level of 0.05 was used.

**Supplemental Figure 3.**
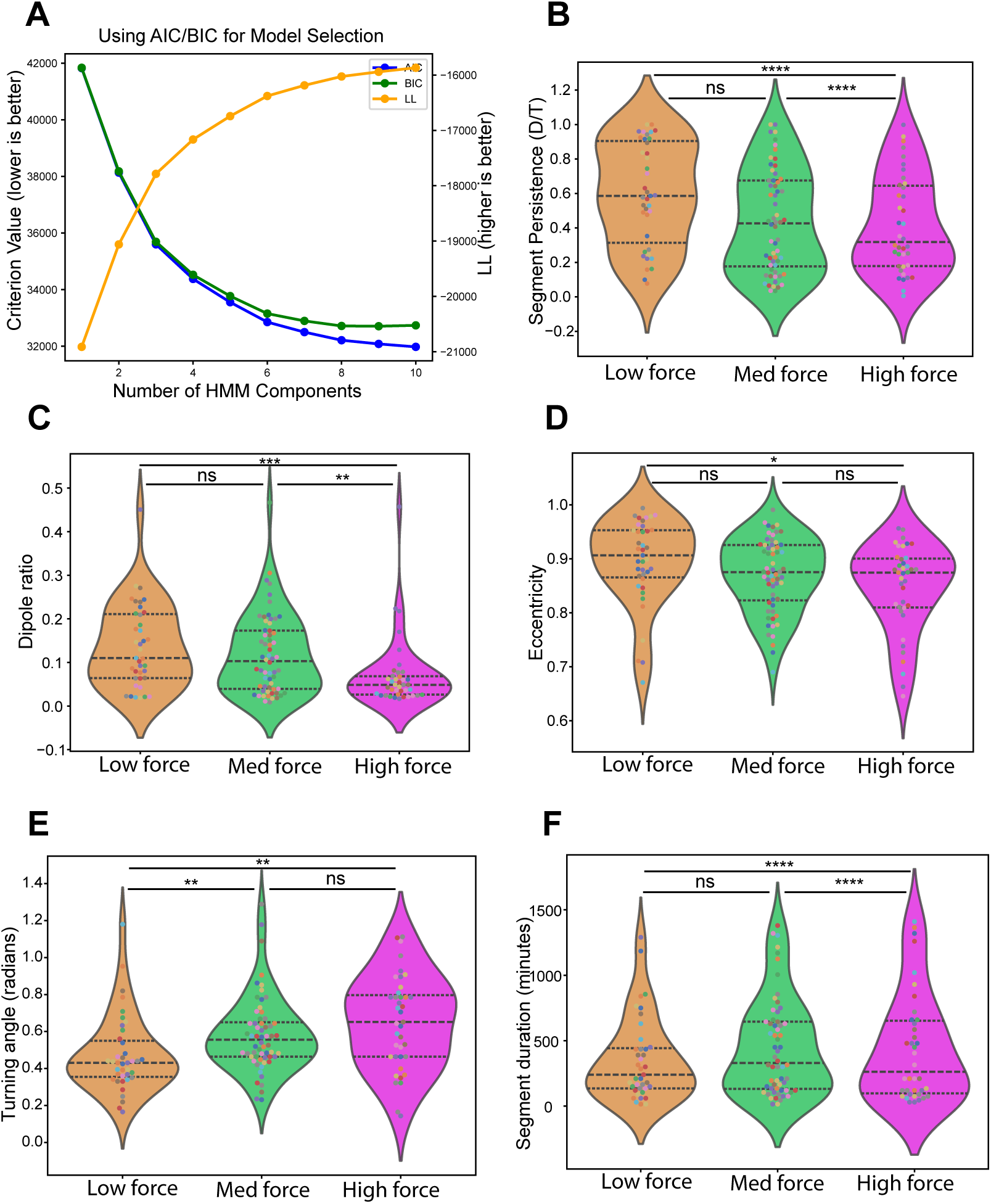
(**A**) AIC (blue), BIC (green), and log-likelihood (yellow) values used to select the appropriate number of states for the Hidden Markov Model for WT cells. (**B-F**) Comparison of median segment values of shape and motion parameters for each HMM state for WT cells. A segment is defined as a series of frames a cell is in one state before it switches states or the track ends. Each dot represents one segment and the colors of the dots correspond to a unique cell (N=52 cells with an average of 2.65 segments per cell). Black dotted lines display the quartiles. Statistical significance is determined with a permutation test comparing the median of the data with the null distribution created from 10,000 permutations with a significance level of 0.05.

**Supplemental Figure 4.**
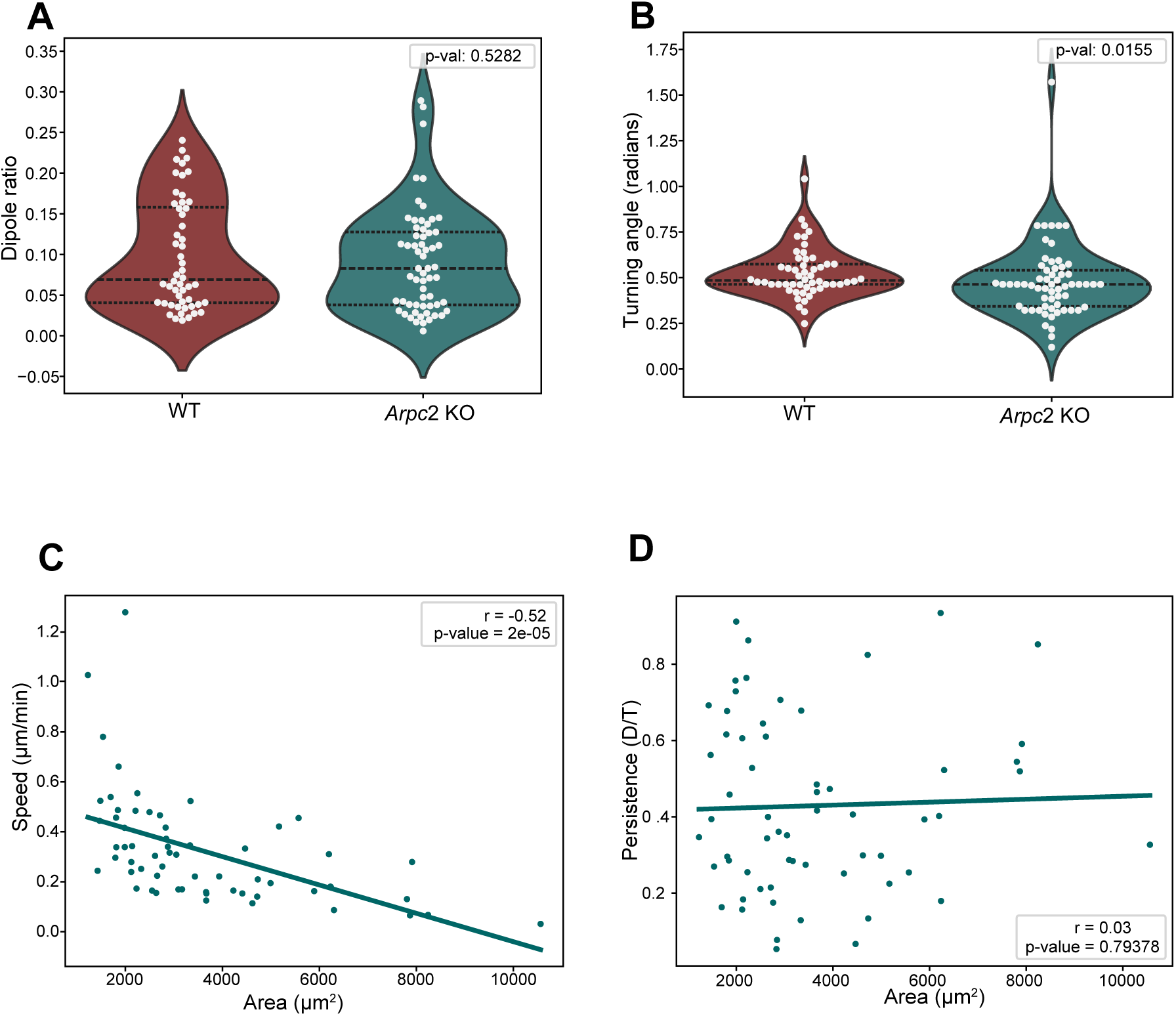
(**A-B**) Comparison of median track values for dipole ratio and turning angle between WT and *Arpc*2 KO cells. Each dot represents the median metric value of one cell over the entire course of its track (WT: N=52 cells, *Arpc*2 KO: N=61 cells). Black dotted lines display the quartiles. (**C-D**) Scatter plots comparing area and motion for *Arpc*2 KO cells. Each point on the plot represents the median value for one cell over the course of its track (N=61 cells). Statistical significance for (**A-B**) is determined with the Mann-Whitney U test. Statistical significance of the Pearson correlation coefficient for (**C-D**) is determined with a two-tailed t-test based on the t-distribution with n-2 degrees of freedom. A significance level of 0.05 was used.

**Supplemental Figure 5.**
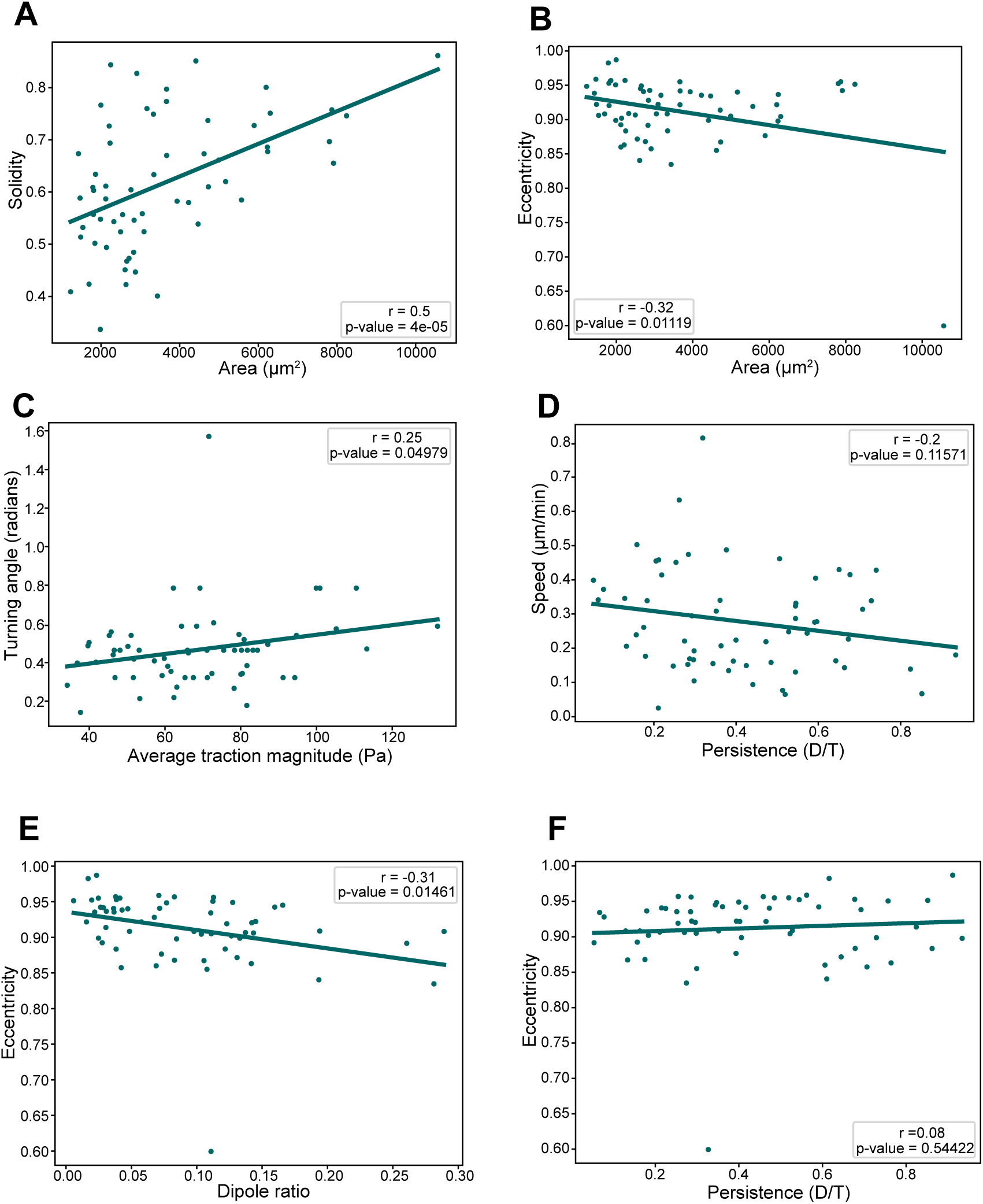
(**A-F**) Scatter plots comparing shape, motion and traction force metrics for *Arpc*2 KO cells. Each point on the plot represents the median value for one cell over the course of its track (N=61 cells). Statistical significance of the Pearson correlation coefficient is determined with a two-tailed t-test based on the t-distribution with n-2 degrees of freedom. A significance level of 0.05 was used.

**Supplemental Figure 6.**
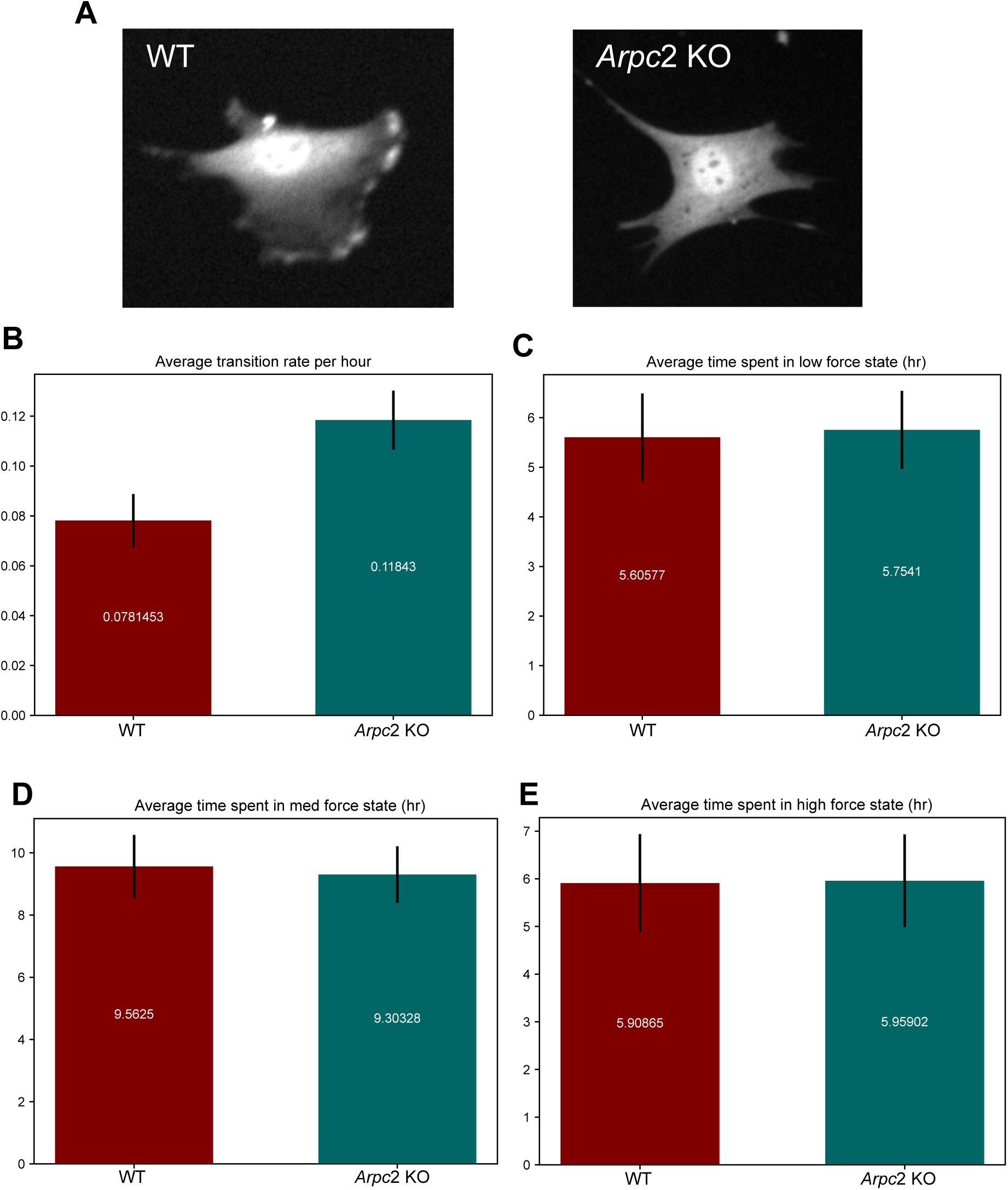
(**A**) Example GFP images of a WT and an *Arpc*2 KO cell. (**B**) Average transition rate per hour for WT and *Arpc*2 KO cells. (**C-E**) Average time spent in each state in hours. (**B-E**) Means are displayed on the bar with the black lines denoting the standard error of mean (SEM).

**Supplemental Figure 7.**
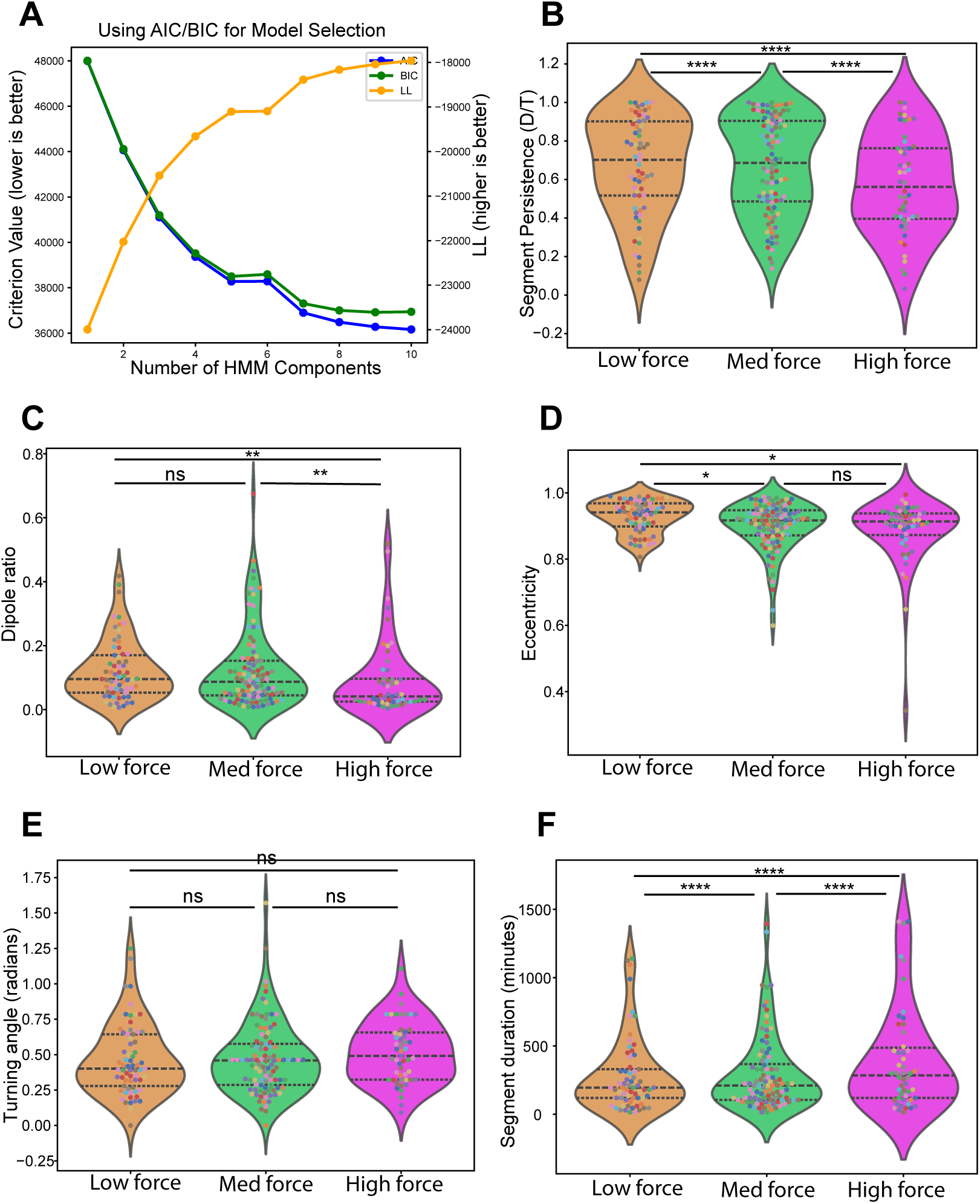
(**A**) AIC (blue), BIC (green), and log-likelihood (yellow) values used to select the appropriate number of states for the Hidden Markov Model for *Arpc*2 KO cells. (**B-F**) Comparison of median segment values of shape and motion parameters for each HMM state for *Arpc*2 KO cells. A segment is defined as a series of frames a cell is in one state before it switches states or the track ends. Each dot represents one segment and the colors of the dots correspond to a unique cell (N=61 cells with an average of 3.49 segments per cell). Black dotted lines display the quartiles. Statistical significance is determined with a permutation test comparing the median of the data with the null distribution created from 10,000 permutations with a significance level of 0.05.

**Supplemental Figure 8.**
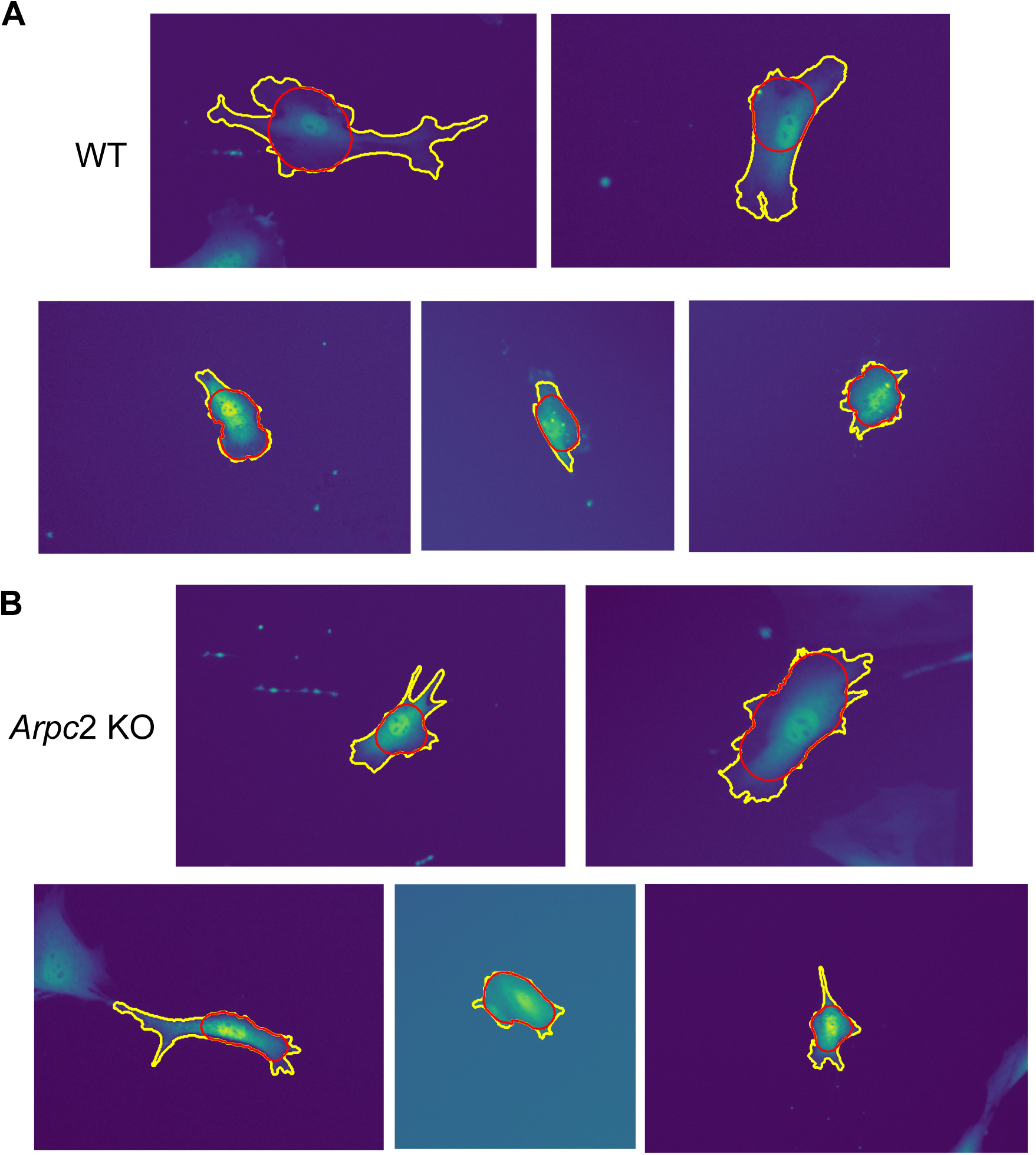
(**A**) WT and (**B**) *Arpc*2 KO cells where the cell mask is outlined in yellow and the cell body is outlined in red. The cell body was obtained from morphological opening (Methods). The nucleus is located within the cell body and everything outside the cell body is referred to as a peripheral protrusion.

**Supplemental Figure 9.**
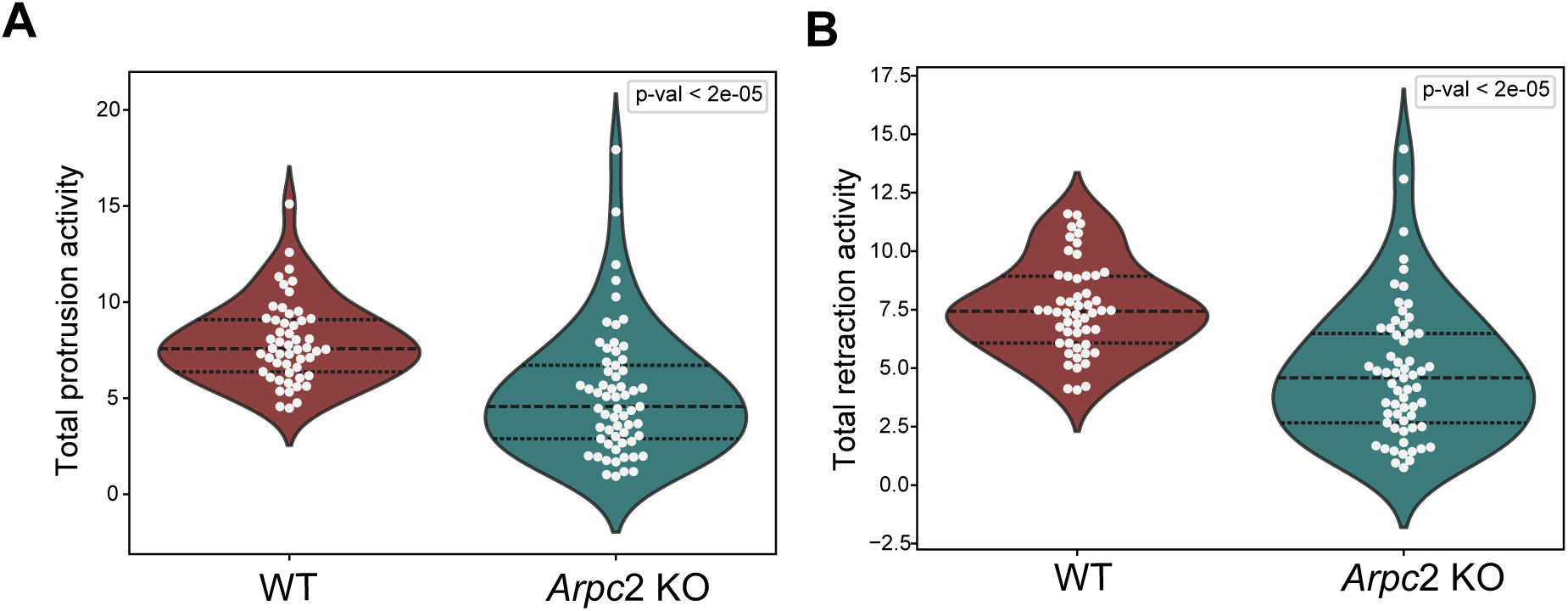
(**A**) Median per cell protrusion boundary activity normalized by cell area and displayed as a percentage. (**B**) Median per cell retraction boundary activity normalized by cell area and displayed as a percentage. WT cells have higher protrusion and retraction boundary activity than *Arpc*2 KO cells. Statistical significance is determined with the Mann-Whitney U test.

**Supplemental Figure 10.**
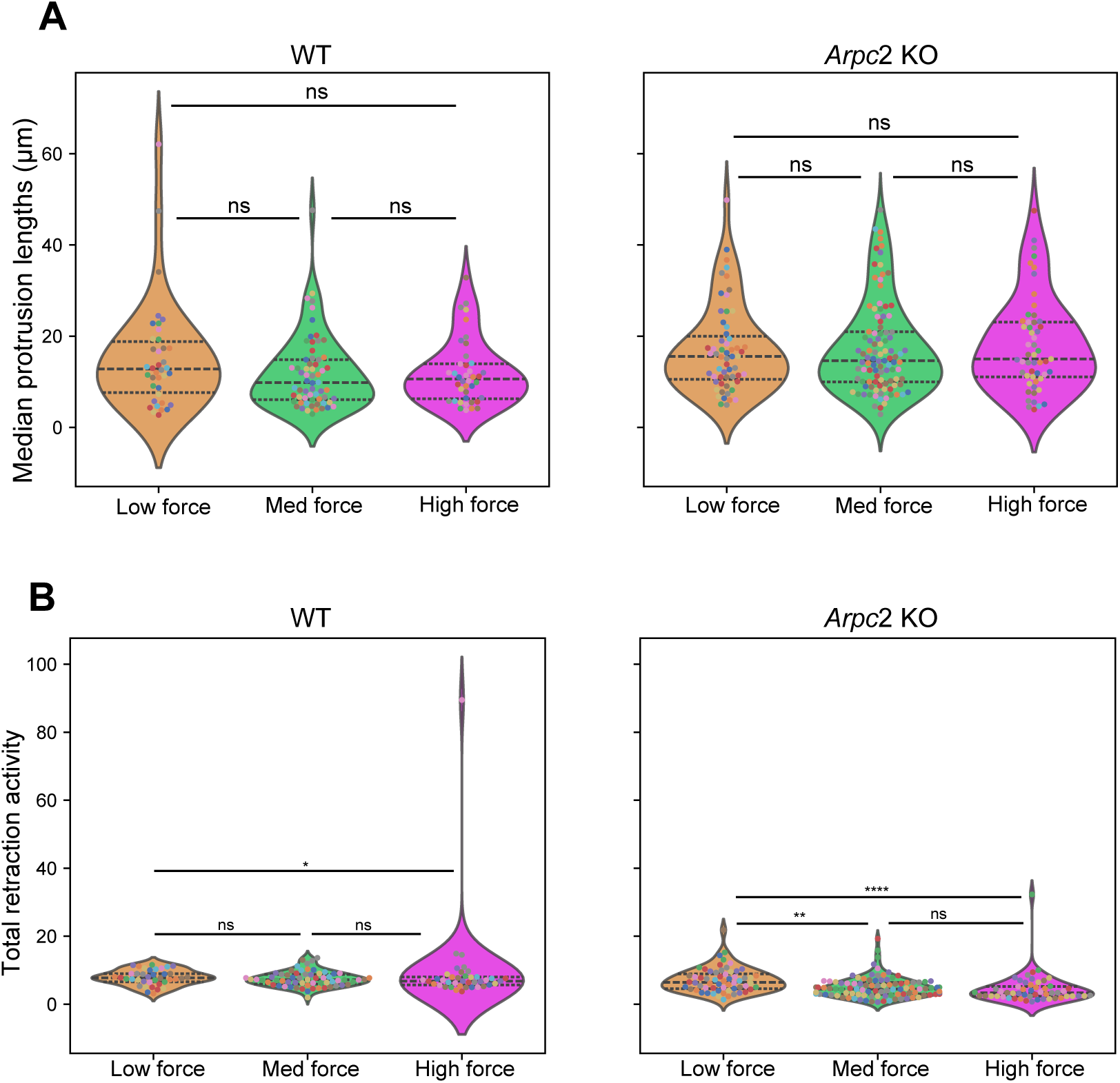
(**A**) The median protrusion length per segment for both WT and *Arpc*2 KO cells in each HMM state. There is no significant difference between the states for either cell type. (**B**) The median retraction activity (normalized by cell area and displayed as a percentage) over a segment within the three predicted hidden Markov model states between WT and *Arpc*2 KO show that the higher force states display slightly lower retraction activity for both WT and *Arpc*2 KO cells. A segment is defined as a series of frames a cell is in one state before it switches states or the track ends. Each dot represents one segment and the colors of the dots correspond to a unique cell. Black dotted lines display the quartiles. Statistical significance is determined with a permutation test comparing the median of the data with the null distribution created from 10,000 permutations with a significance level of 0.05. (WT: N=52 cells, *Arpc*2 KO: N=61 cells)

